# The IKC-leptin axis functions as a molecular bridge between neurodevelopment and endocrine signaling

**DOI:** 10.64898/2026.01.05.697408

**Authors:** Elena Forzisi, Anika Gaur, Aniket Gaur, Maria del Carmen Medranda, Rachel Philip, Srishti Anand, Clotilde Impaloni, Kiarah Leonard, Srinidi Venkateswaran, Federico Sesti

**Author notes:** correspondence to: Federico Sesti web site: http://sestilab.com.

## Abstract

Kcnb1, a voltage-gated K⁺ channel widely expressed in brain, assembles with integrins to form integrin-K⁺ channel complexes (IKCs) in cortex and hypothalamus. Pathogenic *KCNB1* variants cause Developmental and Epileptic Encephalopathy (DEE), and Kcnb1-null (Null) mice reproduce core neurodevelopmental defects alongside chronic hypoleptinemia, suggesting a leptin-IKC signaling axis. We find that Kcnb1, integrins, and the leptin receptor (LepR) co-immunoprecipitate and co-localize in cortical neurons. Perinatal leptin supplementation in Null mice improves cognition, corrects cortical anatomy, and restores neuronal migration, dendritic arborization, and synaptic functionality. In primary Null cortical neurons, leptin normalizes dendritic growth and actin remodeling via integrin-dependent mechanisms. By contrast, leptin has modest effects in WT animals and neurons; sensitivity is unmasked by pharmacological manipulation of integrins, and, strikingly, LepR blockade in the absence of exogenous leptin is sufficient to induce Null-like defects in WT neurons. These findings argue that hypoleptinemia alone does not drive the Null phenotype.

Instead, they reveal a previously unrecognized, dual-mode model in which LepR signaling is regulated by both its ligand and IKCs. This conceptual advance links IKC-LepR coupling to neurodevelopment and DEE and pinpoints their interface as a tractable therapeutic target.

## INTRODUCTION

Leptin is a key hormone regulating energy homeostasis, primarily through its action on hypothalamic neurons in the arcuate nucleus (ARH) that control feeding behavior (1–3). However, the widespread expression of leptin receptors (LepRs) across the brain suggests that leptin also plays broader roles in neural development and function (4).

Consistent with this, disruptions in leptin and insulin signaling are common in maternal obesity, a recognized risk factor for impaired neurodevelopmental and cognitive outcomes in offspring (5,6). Clinical evidence further supports leptin’s involvement: low leptin levels in preterm infants correlate with impaired brain growth, whereas elevated leptin levels are observed in childhood neurodevelopmental disorders such as autism spectrum disorder (ASD) and Rett syndrome (7–10). Animal studies have reinforced leptin’s critical function during brain development, although the molecular pathways remain incompletely defined (11–16). Recent work has implicated the voltage-gated potassium channel Kcnb1 as a mediator of leptin signaling (17). Mice lacking functional Kcnb1 channels (Null mice) display reduced adiposity, constitutive hypoleptinemia, and resistance to leptin’s anorexigenic effects. Importantly, both Null mice and those expressing the pathogenic Kcnb1 R312H variant identified in children with Developmental and Epileptic Encephalopathy (DEE) exhibit severe neurodevelopmental abnormalities, including defective glutamatergic neuron migration, neuromorphological deficits, disrupted axon guidance, and cognitive impairment (18–20). Strikingly, comparable developmental defects occur whether Kcnb1 currents are absent (Null) or preserved (R312H), indicating that Kcnb1’s developmental roles are mediated primarily through non-ionic mechanisms (18). These mechanisms are thought to derive from Kcnb1’s interaction with integrins in Integrin–K⁺ Channel Complexes (IKCs), which regulate actin cytoskeletal remodeling and related neurodevelopmental processes (21,22). Together, these observations suggest the existence of a leptin–IKC signaling axis that integrates metabolic and neurodevelopmental pathways. Here, we test this hypothesis and provide evidence that suggests that constitutive hypoleptinemia resulting from hypothalamic IKC disruption in the Null, may not fully account for the neurodevelopmental impairments observed in these mice. Rather, our results suggest that Kcnb1, integrins, and LepR assemble into a tri-partite complex in the cortex, within which LepR can be modulated by both its cognate ligand (leptin) and IKC-dependent control. If confirmed, this framework would reveal a previously unrecognized mode of leptin signaling in brain development and define a novel interface linking ion channels, integrins, and metabolic hormones.

## RESULTS

### Leptin corrects cognitive deficit in Null mice

Null mice are markedly hyperactive and display broad deficits across sensory processing, executive function, and attention (18). To test whether constitutive hypoleptinemia contributes to these phenotypes, we employed two complementary strategies to enhance early-life leptin exposure: (1) cross-fostering, in which Null pups were reared by WT dams (and *vice versa*), thereby altering leptin levels via maternal milk; and (2) daily intraperitoneal leptin injections (0.5 mg/kg body weight. The surge of blood leptin in 4 week-old animals 1.5 hours after injection is shown in **Fig. S1**) allowing us to isolate leptin-specific effects from maternal or environmental influences. Body weight was monitored throughout treatment to detect potential anorexigenic effects. WT pups showed a modest but significant reduction at 4 weeks (10%; **Fig. S1A**), whereas Null pups exhibited no abnormalities, consistent with their desensitization to leptin’s anorexigenic action. Leptin levels measured at four weeks and eight weeks of age, are shown in **Fig. S1B-D** (as blood sampling is technically challenging in pups we used young animals). Both cross-fostering and exogenous leptin administration restored circulating leptin to normal levels in Null mice at 4 weeks. Leptin levels were also normal in WT mice at this age. In contrast, by 8 weeks WT pups showed a modest trend toward reduced leptin levels, potentially reflecting long-term consequences of their weight loss, while Null animals remained hypoleptinemic. Behavioral testing was performed at 4 weeks at the end of leptin treatment. Hyperactivity, a hallmark phenotype of Null and other Kcnb1-DEE mouse models, resembling ADHD-like traits, was evaluated in the open field (18,23). Cross-fostering or leptin administration had no effect on WT behavior but significantly reduced hyperactivity in Null mice (**Figs. 1A-B**). Interestingly, this effect was sex-specific: leptin markedly attenuated hyperactivity in males but was largely ineffective in females. Next, motivation and stress-coping behavior were assessed with the nest-building assay (**Fig. 1C**). Null mice performed significantly worse than WT controls, showing a striking lack of motivation and organized thinking. Leptin treatment did not affect WT performance but significantly improved nest building in Null mice. Finally, sensorimotor reactivity was tested using the marble-burying paradigm, in which untreated DEE mice typically ignore marbles, a phenotype reminiscent of ASD-like hypo-responsiveness (**Fig. 1D**) (24). As expected, Null mice exhibited little interest in the marbles, whereas WT behavior was unaffected by leptin treatment. Notably, cross-fostering improved marble-burying performance in Null mice of both sexes, while leptin injections had minimal effect, suggesting a strong maternal contribution to this behavior. Together, these results demonstrate that perinatal leptin oscillations are unlikely to contribute significantly to cognitive function in WT mice. By contrast, they demonstrate that early-life leptin supplementation can ameliorate several cognitive and behavioral deficits in Null mice. Importantly, the effects were selective, showing sex dependence in hyperactivity and a strong maternal contribution to sensorimotor responsiveness.

**Figure 1.**
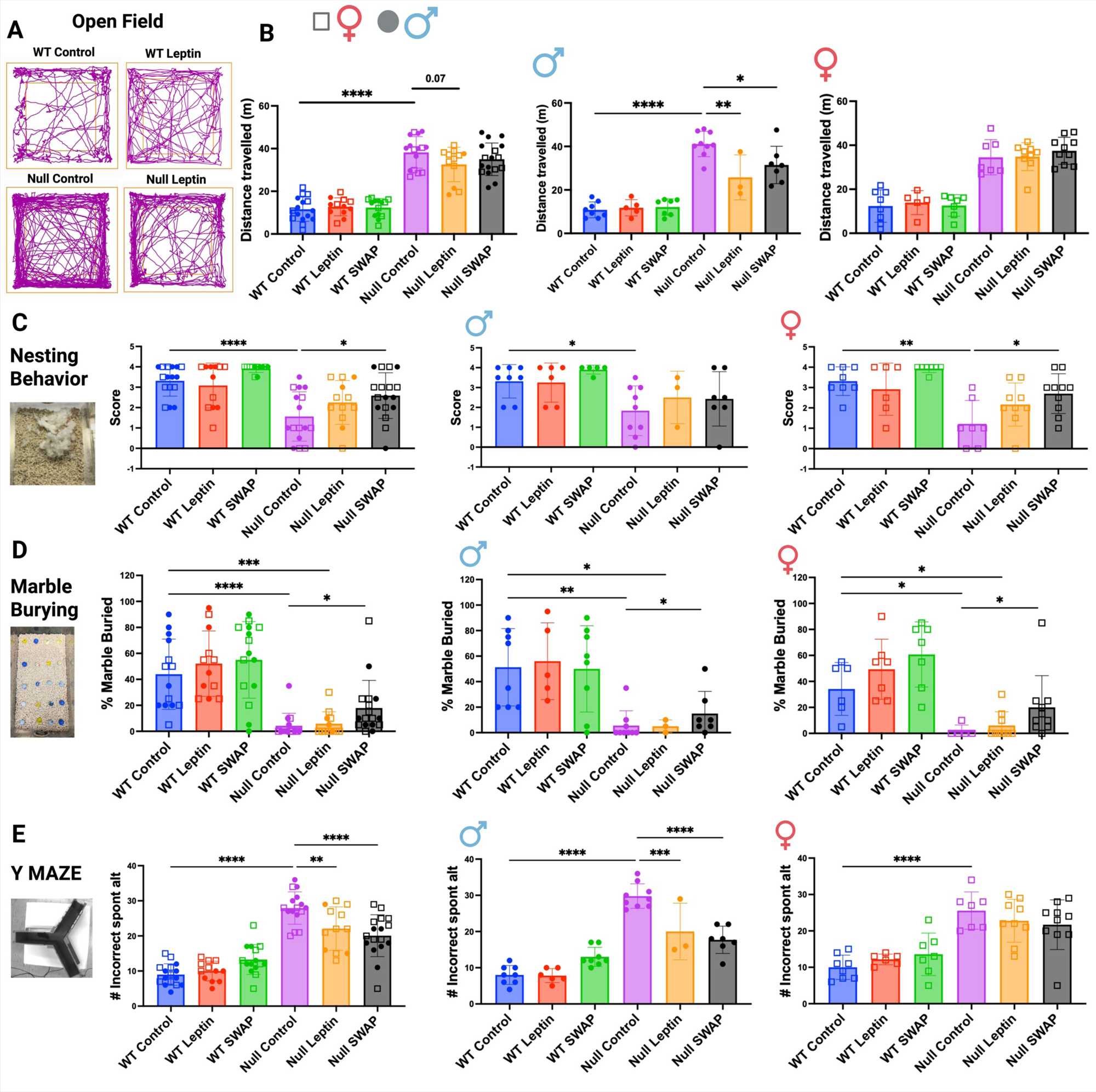
Leptin improves cognitive function in Null mice. A) Representative walking plots of WT and Null mice in the open field arena, reared in control or subject to leptin injections. B) Average distance run in the open field arena for WT and Null mice reared in the indicated experimental conditions. C) Nesting score (see methods) for WT and Null mice reared in the indicated experimental conditions. D) Percentage of marbles buried by WT and Null mice reared in the indicated experimental conditions. E) Number of incorrect spontaneous alternations in the Y-maze arena for WT and Null reared in the indicated experimental conditions. Four-week-old male and female mice received daily injections of saline (control), or 0.5 mg/body weight recombinant leptin (leptin) from P0–P28, or were cross-fostered (SWAP), as described elsewhere. Behavioral testing was performed using an EthoVision tracking system and analyzed with ANY Maze software. Data are shown both as totals and stratified by sex. Statistical comparisons were performed using one-way ANOVA with Tukey’s post hoc. N=12-16 mice/condition.

### Leptin has mixed effects on learning and memory in mice

Given clinical evidence linking leptin deficiency to impaired learning in children, we assessed cognitive performance in three assays: the Spontaneous Alternation Y-maze (spatial working memory), the Novel Object Recognition (NOR) test (short-term memory), and the Barnes maze (spatial learning and memory). In the Y-maze, Null mice performed significantly worse than WT controls (**Fig. 1E**). Leptin administration did not alter WT performance but significantly improved Null performance, though not to WT levels. The effect was sex-dependent, with robust improvement in males and only a modest trend in females. In the NOR test, Null mice once more showed significantly poorer performance than WT, despite similar engagement with the objects across all groups (**Fig. S2**).

Leptin treatment significantly improved object recognition in Null animals, with injections producing a greater effect than cross-fostering. WT performance remained unchanged. In the Barnes maze, WT and Null mice performed comparably during the acquisition phase, as measured by the time to locate and enter the escape hole (**Fig. S3A-F**).

Leptin enhanced exploratory behavior in Null mice, such that after learning the escape location by day 3, they continued to explore the maze rather than proceeding directly to the target (**Fig. S3F**). During the retention phase, Null mice performed modestly worse than WT (P=0.0505), and leptin increased escape hole entries to a similar extent in both genotypes (**Fig. S3G-H**). Additionally, leptin-treated Null mice spent more time in the quadrant housing the escape hole during the acquisition phase, potentially underscoring subtle enhancement of spatial memory (**Fig. S3I-J**). Together, these results demonstrate that while hippocampal-dependent spatial learning and memory are preserved in Null mice, leptin supplementation selectively improves working memory and object recognition. Overall, leptin’s mixed and context-dependent effects underscore the complexity of its actions on cognitive function.

### Leptin corrects neuroanatomical defects in Null mice

Behavioral impairments in Null mice are accompanied by widespread neuroanatomical abnormalities that likely contribute to their cognitive deficits (24). Key structural defects include reduced thickness of upper cortical layers (LII/III), as well as anomalies in other regions such as the *corpus callosum* (CC). To assess whether leptin can ameliorate these abnormalities, we examined brain sections stained with neurofilament medium (NFM), L1CAM, and microtubule-associated protein 2 (Map2). Representative NFM immunofluorescence images of coronal sections from untreated and leptin-treated mice are shown in **Figs. 2A-B**, with corresponding quantifications of cortical layer thickness presented in **Fig. 2C**. These results revealed that LII/III/IV are significantly thinner in Null mice compared to WT, whereas LV/VI are comparable, consistent with prior observations with another mouse model of DEE (24). Strikingly, while perinatal leptin supplementation had no detectable effect on WT cortical layers, it restored normal layers thickness in the Null.

**Figure 2.**
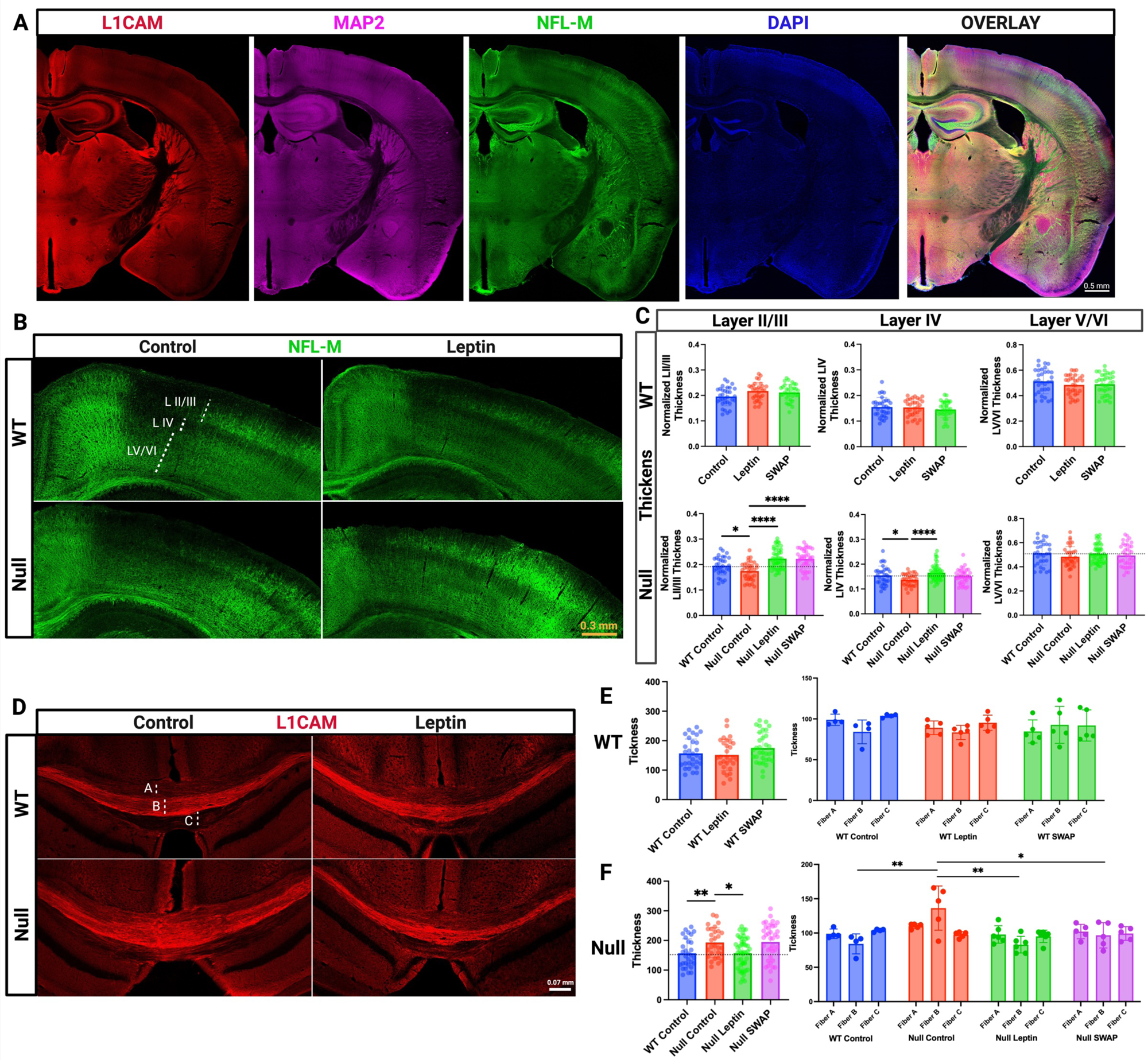
Leptin improves cortical neuroanatomy in Null mice. A) Representative confocal images of a neocortical section from WT, co-immunostained for L1CAM (red), Map2 (magenta), NFM (green) with DAPI counterstain (blue). B) Representative confocal images of neocortical sections from WT and Null mice reared under the indicated conditions, immunostained for NFM (green). Cortical layers are indicated. C) Quantifications of cortical layer thickness normalized to total thickness, based on NFM immunofluorescence, as shown in (B). D) Representative confocal images of the corpus callosum from WT and Null mice under the indicated conditions, immunostained for L1CAM. E) Quantification of corpus callosum fibers thickness, in WT and Null mice across experimental groups. Data are shown as totals and stratified by fiber. Eight-week-old male and female mice received daily injections of saline (control), or 0.5 mg/body weight recombinant leptin (leptin) from P0–P28, or were cross-fostered (SWAP), as described elsewhere. N=3-5 brains/condition with 5-7 technical replicates/brain (points in graphs). Statistical comparisons were performed using one-way ANOVA with Tukey’s post hoc test.

The effects of leptin on the *corpus callosum* visualized by L1CAM immunofluorescence, are shown in **Figs. 2D-F**. Null mice exhibit abnormally thick *corpi callosi*, particularly the commissural projections of the *splenum* (Fiber B, **Fig. 2F**). Leptin did not alter CC morphology in WT mice, but it fully normalized CC anatomy in the Null. In summary, exogenous leptin administration rescues both cortical and subcortical neuroanatomical defects in Null mice, underscoring its potential to correct fundamental developmental abnormalities. In contrast, as seen at behavioral level, leptin has no impact on cortical and subcortical neuroanatomical features of WT brains.

### Leptin rescues neuronal migration defects in Null mice

Null embryos show severe impairments in the radial migration of glutamatergic neurons destined for the upper cortical layers (UL), likely contributing to cortical malformations (18). To test whether leptin supplementation restores migration, pregnant dams received 5-bromodeoxyuridine (BrdU) at embryonic day 16.5 (E16.5) to label UL-bound neurons. In a parallel regimen, dams (from E16.5) and then offspring (through postnatal day 7, P7) were injected with leptin as described. These neurons are born in the ventricular zone (VZ) and normally traverse deep layers (DL) en route to the UL. At P7, when migration is complete, brains were harvested and BrdU+ cells quantified. As expected, Null mice exhibited pronounced migration deficits, with BrdU+ progeny abnormally retained in the DL and a consequent altered UL distribution (**Fig. 3**) (18). Notably, perinatal leptin largely normalized migration in Null mice, leaving only few BrdU+ cells in the DL and correcting UL lamination. In contrast, leptin produced no effect in WT. Thus, perinatal leptin supplementation rescues defective neuronal migration in Null mice without perturbing migration in WT. These data are consistent with, and may help explain, earlier embryonic studies in which leptin supplementation in constitutively hypoleptinemic ob/ob embryos promoted migration of neuronal lineage cells to the cortical plate and reshaped cortical neuronal gene expression programs, supporting a role for leptin in corticogenesis (25,26).

**Figure 3.**
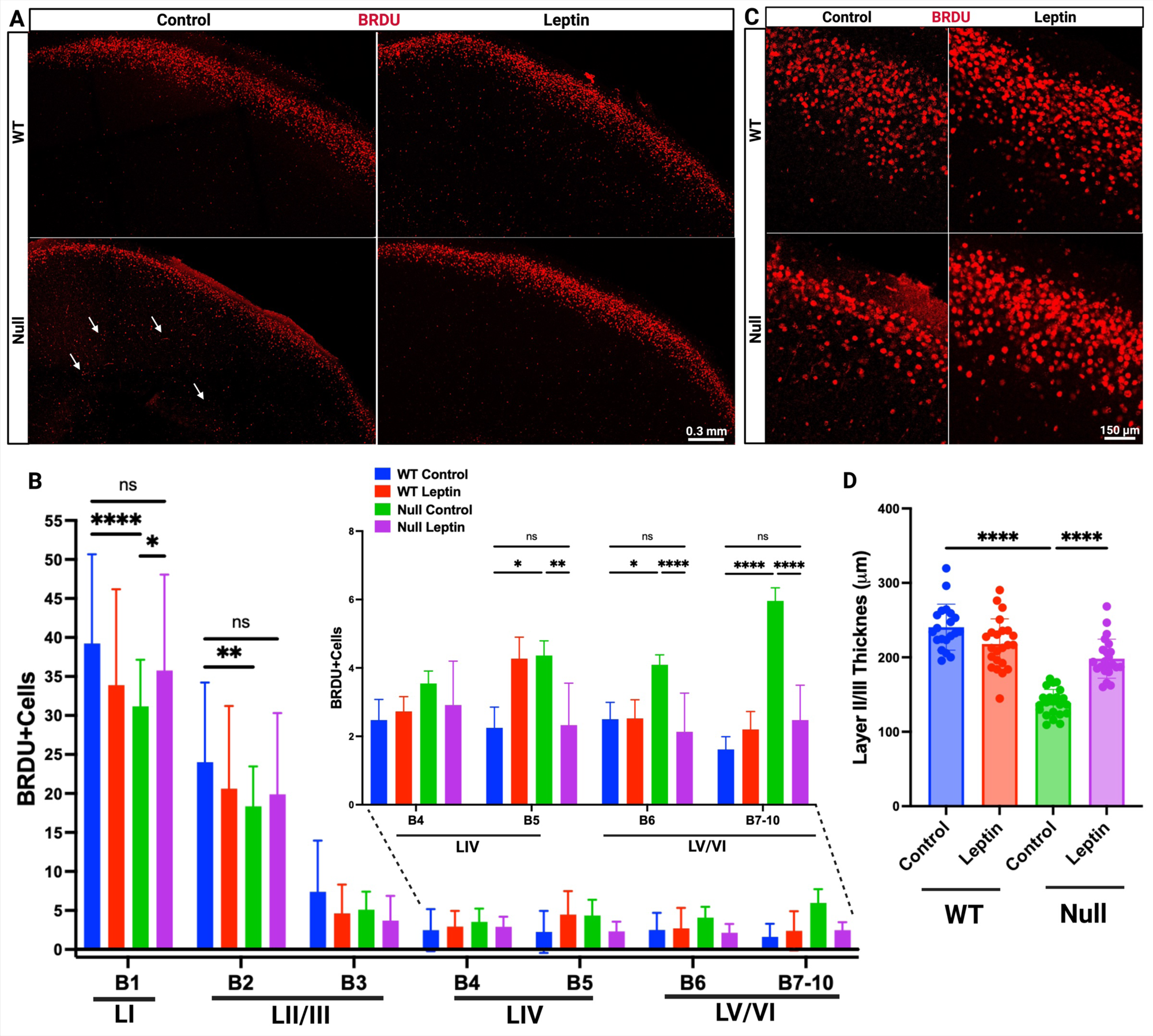
Leptin corrects migratory defects in the Null cortex. A) Representative confocal images of neocortex stained for BrdU. Examples of BrdU⁺ cells retained in deep layers (IV–VI) are indicated by arrows. B) Quantification of BrdU⁺ cell distribution across cortical layers. The cortical plate was divided into 10 equal bins, and the relative distribution of BrdU⁺ cells per bin was plotted. *Inset*: magnification of B4-B10 distribution. Correspondence between bins and cortical layers is indicated. C) Representative confocal magnifications of images in (A), showing BrdU^+^ cells distribution in LII/III. D) Quantifications of LII/III thickness for WT and Null under the indicated conditions. BrdU was administered at E16.5, when upper-layer neurons are born. Analyses were performed at P7. N = 3 brains per condition; for each brain, 6-8 coronal sections were examined with two technical replicates per section. Statistical comparisons were performed using one-way ANOVA with Tukey’s post hoc test.

### Leptin enhances synaptic functionality in Null mice

A hallmark of Null brains is impaired synaptic function, reflected by fewer co-localizations between the presynaptic marker synapsin-1 (Syn-1) and the postsynaptic marker PSD-95 (18). Consistent with this, immunofluorescence analyses showed reduced PSD-95/Syn-1 co-localization in Null versus WT cortices (**Fig. S4**). Leptin treatment significantly increased PSD-95/Syn-1 co-localization in Null mice, restoring it to WT levels. In WT animals, cross-fostering had no effect, whereas leptin injections decreased co-localization. Thus, leptin rescues a synaptic apposition deficit in Null mice, while producing a context-dependent reduction in synaptic co-localization in WT.

### Leptin influences cortical neurons development

To investigate the interplay between leptin and Kcnb1 in cortical neurodevelopment, we examined neuronal morphology, a process strongly regulated by Kcnb1, using complementary *in vivo* and *in vitro* approaches. Golgi staining of cortical tissue allowed structural assessment *in situ*, while primary neuronal cultures provided a system amenable to pharmacological and genetic manipulation, thereby enabling us to establish molecular correlations between the channel and the hormone. Qualitative inspection of Golgi-impregnated cortices revealed aberrant dendritic arborization in Null mice compared with WT, consistent with prior observations (**Figs. 4A-B**) (18). To quantify these abnormalities, we conducted Sholl analysis, assessing dendritic complexity as the Sholl Intersection Profile (SIP) normalized to cell number within the area of maximum radius *r* (150 μm, example in the inset of **Fig. 4A**), and measuring total dendritic length within a 100 µm square containing no more than a single soma (dendritic density). In LII/III, Null neurons displayed significantly greater dendritic intersections than WT. Leptin treatment had no effect on WT but significantly reduced dendritic complexity in Null, normalizing it to WT levels (**Figs. 4C-D**). A similar pattern was observed in LV/VI, though baseline differences were smaller (**Figs. 4C-D**). Leptin likewise normalized dendritic density in Null neurons across both LII/III and LV/VI, while leaving WT unaffected (**Fig. 4E**). These findings at first appear inconsistent with **Fig. 2C**, which shows reduced thickness of LII/III in Null cortex, rescued by leptin as measured by NFM immunostaining. However, dendritic complexity (Golgi analysis) and cortical layer thickness (immunostaining) capture distinct features of neuronal architecture. NFM staining primarily highlights axonal architecture, providing an indication of how neurons organize to form cortical layers. By contrast, an increase in dendritic material, whether through additional branching or elongation, would generally be accompanied by elevated expression of NFM. Indeed, fluorescence intensity normalized across preparations revealed higher NFM levels in Null versus WT, with leptin reducing these signals to WT values, again without effect in WT mice (**Fig. S5**).

**Figure 4.**
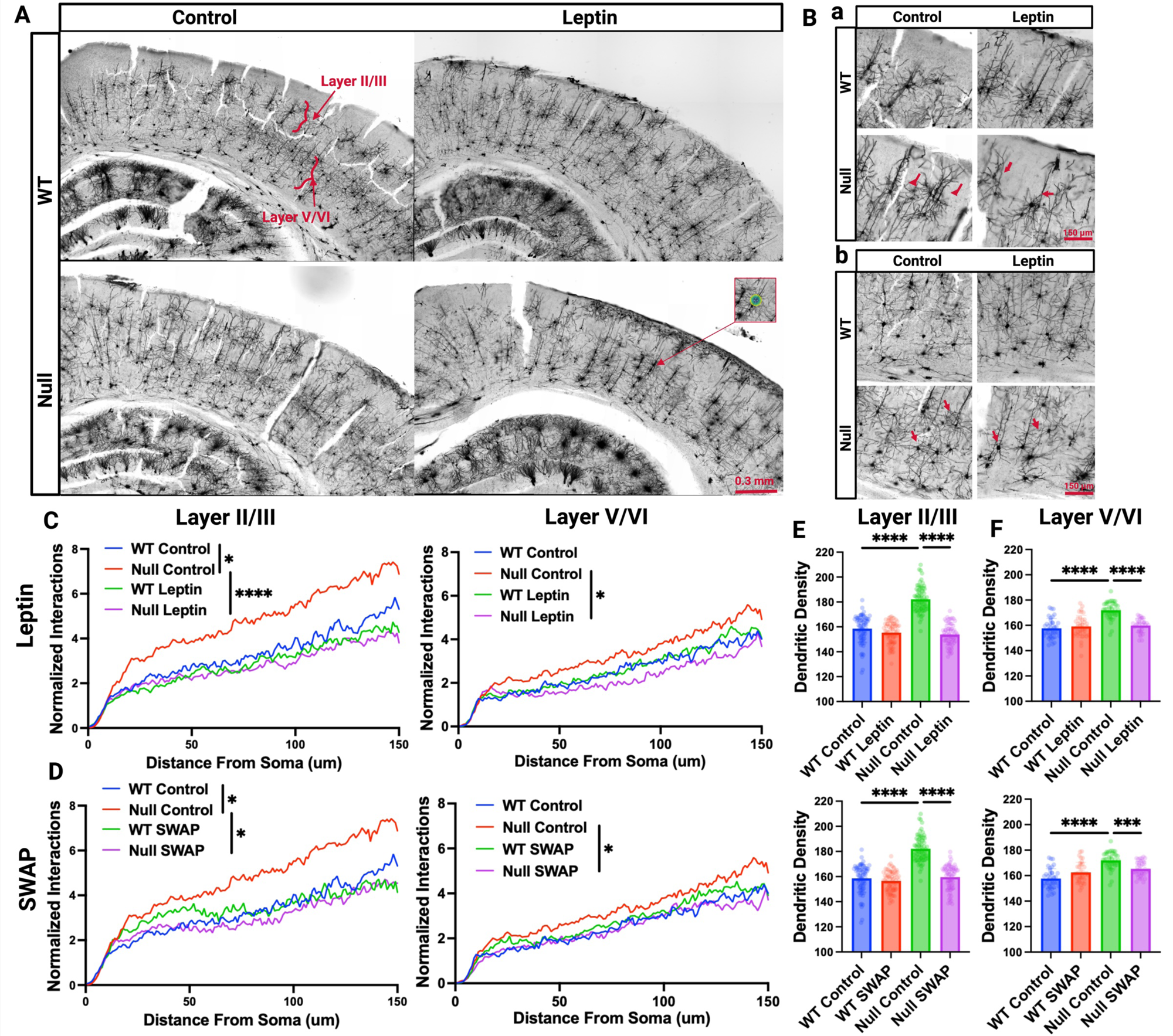
Leptin corrects dendritic over-arborization in Null mice. A) Representative Golgi-stained cortical sections from WT and Null mice under the indicated conditions. *Inset*: representative Sholl analysis. B) Higher-magnification images from (A) highlighting dendritic over-arborization. a) Layer II/III b) Layer V/VI. C) Sholl analyses of average number of dendritic intersections in layers II/III and layers V/VI as a function of the distance from a neuronal soma, for pups injected with leptin. Intersections were normalized by the number of cells within a circular area of r=150 μm (example inset in (A)). D) As in (C), for pups cross-fostered (SWAP). In (C) and (D) each curve was calculated from S≥50-80 Sholl analyses. E) Dendritic density, quantified as total dendritic length within a 100 µm × 100 µm square containing no more than one soma, in layers II/III (left column) and V/VI (right column) under the indicated conditions. Brains of four-week-old mice treated with daily injections of saline (control), or 0.5 mg/kg body weight recombinant leptin from P0–P28 (leptin), or cross-fostered (SWAP), were processed with Golgi staining and analyzed by confocal microscopy. For each brain (N = 3-7 per condition), 8–10 images were acquired. Statistical comparisons were made using one-way ANOVA with Tukey’s post hoc test.

### Leptin increases dendritic spine maturation in the Null

Spine morphogenesis was quantified on Golgi–silver–impregnated sections (**Fig. 5A**). As reported, Null dendrites bear significantly fewer spines than WT (**Fig. 5B**) (18). Perinatal leptin did not change total spine number in WT, but it restored it in Null to WT levels. Class-distribution analyses point to a leptin-driven neurotrophic effect in Null neurons: leptin increased mushroom spines with a reciprocal decrease in immature (thin and filopodia) spines, while stubby spines were largely unchanged (**Fig. 5C**). The pie charts in **Fig. 5D** further underscore genotype specificity. In Null, leptin consistently shifted the population toward the mushroom class at the expense of immature spines. In WT, the mushroom fraction was essentially stable across treatments, but the other classes redistributed: leptin increased immature spines (thin ∼ +10%; filopodia ∼ +60%) predominantly at the expense of premature stubby spines (**Fig. S6** and **Fig. 5D**). This pattern aligns with the reduced synaptic co-localization observed in leptin-treated WT cortices (**Fig. S4**).

**Figure 5.**
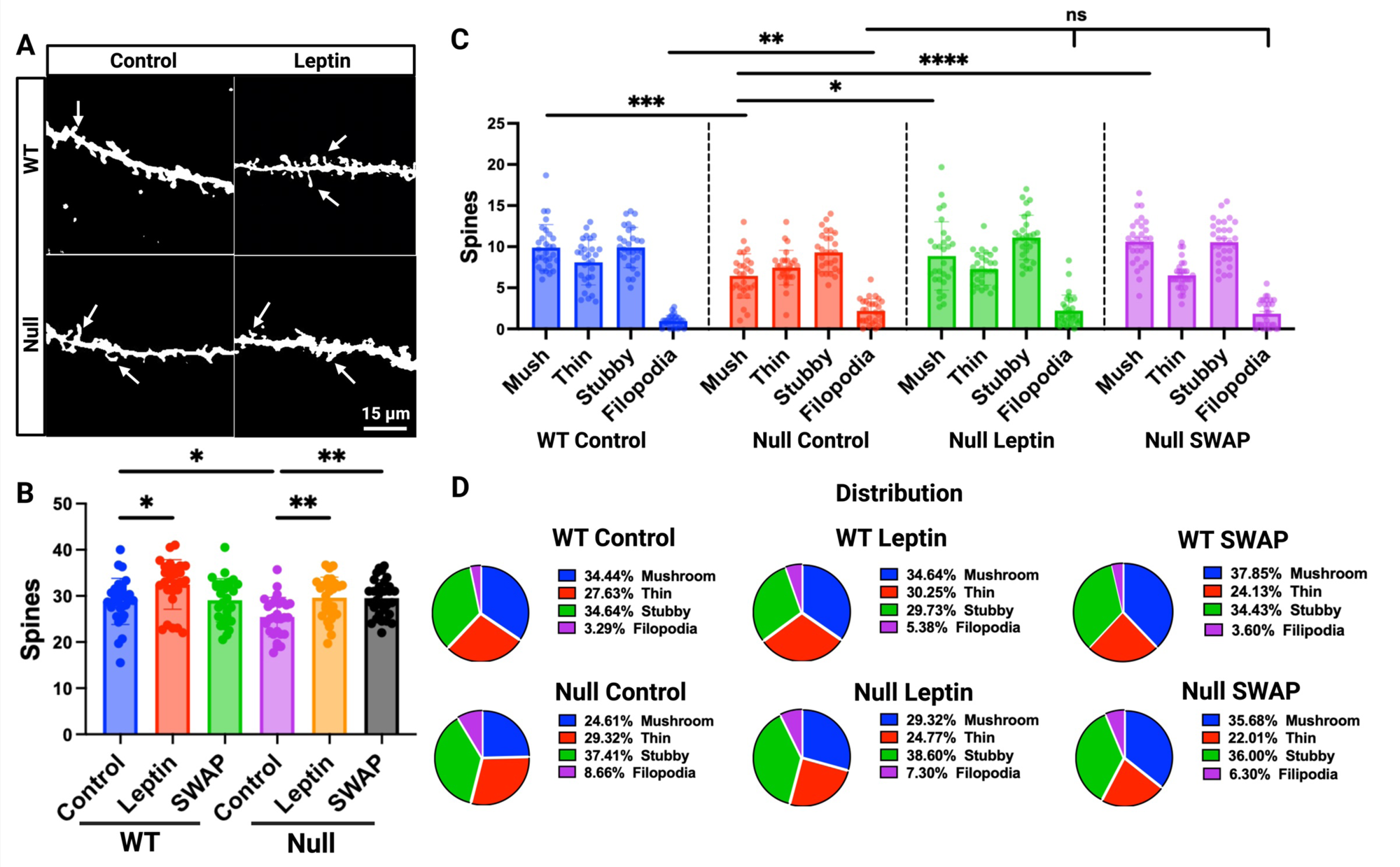
Leptin corrects spine morphology in Null mice. A) Representative confocal images of cortical dendrites from WT and Null mice under the indicated conditions. B) Quantification of total dendritic spine number across conditions. C) Distribution of spine morphologies (mushroom, thin, stubby, filopodia) under each condition. D) Pie charts illustrating the proportion of spine classes for each genotype and treatment. Brains of four-week-old mice treated with daily injections of saline (control), or 0.5 mg/body weight recombinant leptin from P0–P28 (leptin), or cross-fostered (SWAP), were processed with Golgi staining and analyzed by confocal microscopy. For each brain (N = 3 per condition), 8–10 images were acquired. Spine counts were performed along fixed dendritic segments (45 µm; 3 dendrites per image) using SpineJ plugin in ImageJ and averaged across individual data points. Statistical comparisons were made using one-way ANOVA with Tukey’s post hoc test.

Collectively, these data indicate that when Kcnb1 function is compromised, leptin promotes both spine formation and maturation, whereas in WT cortex it subtly biases the spine landscape toward immaturity, consistent with a Kcnb1-dependent tuning of leptin’s neurotrophic effects.

### Leptin affects the neural architecture of cultured neurons

Primary cortical neurons were analyzed as in **Fig. 6**. Our readouts included neuronal morphology (to align *in vivo* with *in vitro* findings) and remodeling of the actin cytoskeleton, a key molecular driver of these morphological processes (28). Dendritic complexity was evaluated using the SIP of individual neurons, along with the length of the apical dendrite (28). At the molecular level, we quantified actin polymerization by FRAP of F-actin in basal dendrites and by measuring the mean size of actin aggregates in the soma and in proximal, large-caliber dendrites (**Fig. 6D**, arrows). These aggregates, which are typically enlarged in DEE neurons, likely reflect a neuron-wide disruption of actin homeostasis. One plausible contributor is altered actin polymerization that shifts the balance between globular (G-) and filamentous F-actin; however, other mechanisms, such as reduced actin turnover, stress-associated cytoskeletal lesions, or sequestration of actin into relatively insoluble inclusions, may also contribute (24,29,30). Notably, the aggregates showed strong co-localization with LepR, detected using a previously validated antibody (WT = 50.2 ± 4.9%; Null = 71.0 ± 4.6%) (17). Consistent with *in vivo* Golgi analyses, Null neurons exhibited marked over-arborization relative to WT (**Fig. 6B**). Leptin had no detectable effect on WT neurons (**Fig. 6C**) but normalized dendritic arborization in Null neurons to WT levels (**Figs. 6D**). Apical dendrites were shorter in WT than in Null; leptin produced opposite effects, increasing apical length in WT while decreasing it in Null (**Figs. 6E-F**). As expected, actin clumps were larger in Null than in WT; leptin increased aggregate size in WT but left it largely unchanged in Null (**Fig. 6G**). For FRAP, F-actin was visualized with Lifeact-GFP (24). Transfection efficiency, evaluated by Lifeact-GFP fluorescence, was comparable across genotypes and conditions (**Fig. S7A**). In this system, fluorescence recovery, R(t) (eqn. 2), is assumed to be reaction-limited, meaning that binding of Lifeact-GFP to F-actin predominates over diffusion into the bleached region (24). Under these conditions, R(t) follows an exponential course determined solely by the unbinding rate constant k_off_ (eqn. 3). Mean R(t) curves for WT and Null in the absence/presence of leptin are illustrated in **Fig. 6H**. Accordingly, fluorescence recovery curves were well fit to a single exponential function, with k_off_ shown in **Fig. 6I**. Null neurons displayed a greater k_off_ than WT (indicating faster recovery), but leptin treatment had no effect on either genotype. We quantified FRAP amplitudes by the mobile fraction (steady-state R(t); eqn. 4), which reflects the exchangeable pool of Lifeact-bound actin and, by extension, the degree of F-actin polymerization dynamics. Relative to WT, Null neurons showed a FRAP signature consistent with actin over-polymerization/dysregulated turnover (**Fig. 6J**). Notably, leptin had little effect in WT but restored the Null mobile fraction to WT levels (**Fig. 6K**). In summary, convergent assays support a coherent model: first, at baseline, Null neurons exhibit pronounced structural and cytoskeletal abnormalities, aberrant dendritic arborization, elongated apical dendrites, and enlarged actin aggregates, together with hyperactive actin polymerization, consistent with the notion that the IKC-leptin axis acts as a brake of these processes. Second, leptin induces only modestly adverse, Null-like, changes in WT neurons (more actin aggregates and longer apical dendrites) but robustly rescues the morphological and cytoskeletal defects in Null neurons, mirroring the *in vivo* phenotype.

**Figure 6.**
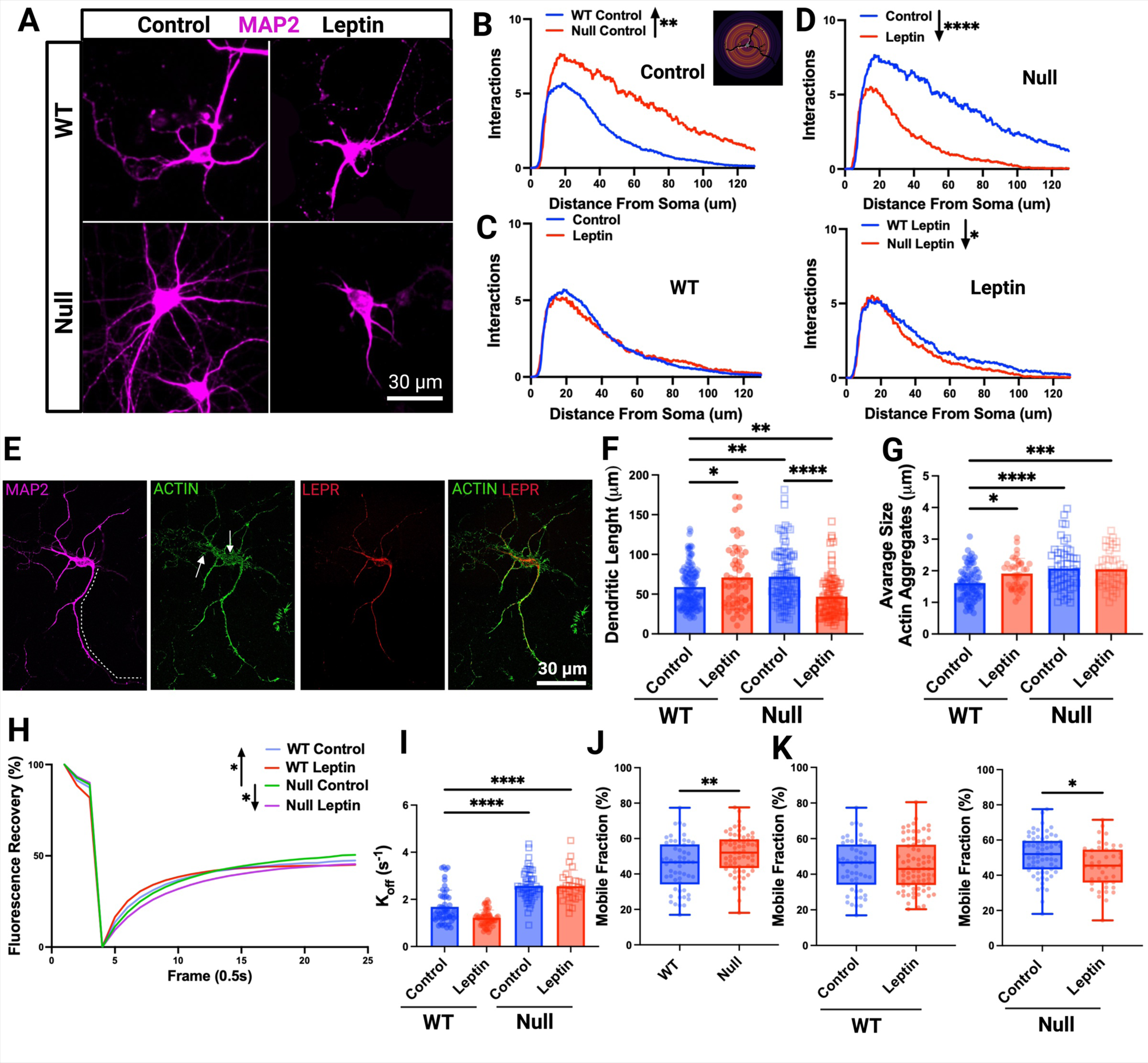
Effects of leptin in primary cortical neurons recapitulate those observed in vivo. A) Representative confocal images of Map2-stained cortical neurons under the indicated conditions. B) Comparisons of the mean SIPs of WT and Null grown in control. *Inset*: representative Sholl analysis. C) Comparisons of the mean SIPs of WT neurons grown in the absence/presence of leptin. D) Comparisons of the mean SIPs of Null neurons grown in the absence/presence of leptin and comparisons of the mean SIPs of WT and Null neurons grown in the presence of leptin. E) Representative confocal images of a cortical neuron co-immunostained for Map2, actin and LepR and overlay of actin and leptin stainings. The dotted line in Map2 shows a reconstruction of the apical dendrite and arrows show actin aggregates. F) Apical dendrite length of WT and Null neurons in the absence/presence of leptin. G) Mean size of actin aggregates for WT and Null neurons in the absence/presence of leptin. H) Fluorescence recovery curves (eqn. 2) for the indicated genotypes and conditions (SEM not shown for clarity). I) Unbinding k_off_ constants, obtained by single exponential fitting of the recovery curves (eqn. 3). J) Comparisons of the mobile fractions (eqn. 4) of WT and Null. K) Mobile fraction control/leptin for WT and Null. For morphological analyses, primary cortical neurons were cultured under control conditions or treated from DIV1 to DIV10 with leptin (0.3 µM). Neurons were fixed and immunostained for Map2 and actin at DIV10, imaged by confocal microscopy, and analyzed using Fiji. N=60-200 neurons genotype/condition from 3-7 biological replicates. For FRAP, neurons were transfected with Lifeact-GFP at DIV7, subjected to photobleaching at DIV8, and fluorescence recovery was quantified. Prior to imaging 1.3 μM leptin was applied for 30 min. N = 45-100 neurons per genotype/condition with 2 technical replicates from 3 biological replicates per genotype/condition. Statistical analyses included the Kolmogorov–Smirnov test for Sholl intersection profiles (SIPs) and one-way ANOVA with Tukey’s post hoc test for dendritic length and mobile fraction.

### Kcnb1, integrins, and LepR form macromolecular complexes

Guided by hypothalamic evidence that LepR associates with Integrin-Kcnb1 complexes, we next tested whether a similar association exists in the cortex (17). Previous studies have shown that *LepR* mRNA trascripts are particualrtly abundant in the upper cortical layers, which are also the layers most affected in the Null (27). Accordingly, RNAscope imaging (**Fig. 7A**) confirmed cortical expression of *Kcnb1* and *LepR* transcripts. In WT animals, *LepR* mRNA displayed a laminar distribution consistent with cortical neuronal organization, whereas this pattern was disrupted in Null animals (**Figs. 7B-C**). Accordingly, imaging of primary cortical neurons revealed extensive co-localization of LepR, Kcnb1 and integrin-β1 within the soma and basal dendrites (**Fig. 7D**-**F**), in line with their broad cortical expression. Five co-immunoprecipitation experiments (representative example in **Fig. 7G**) further demonstrated that in WT cortical lysates, LepR co-precipitates with Kcnb1 and integrins β1 and β5. As expected, Kcnb1 was absent from LepR pulldowns in Null lysates, although LepR continued to associate with integrins. Together, these results establish that LepR physically interacts with Kcnb1 and integrins in the cortex. While the precise molecular architecture and stoichiometry remain to be elucidated, the data strongly support a model in which LepR, Kcnb1, and integrins co-assemble into a shared macromolecular complex.

**Figure 7.**
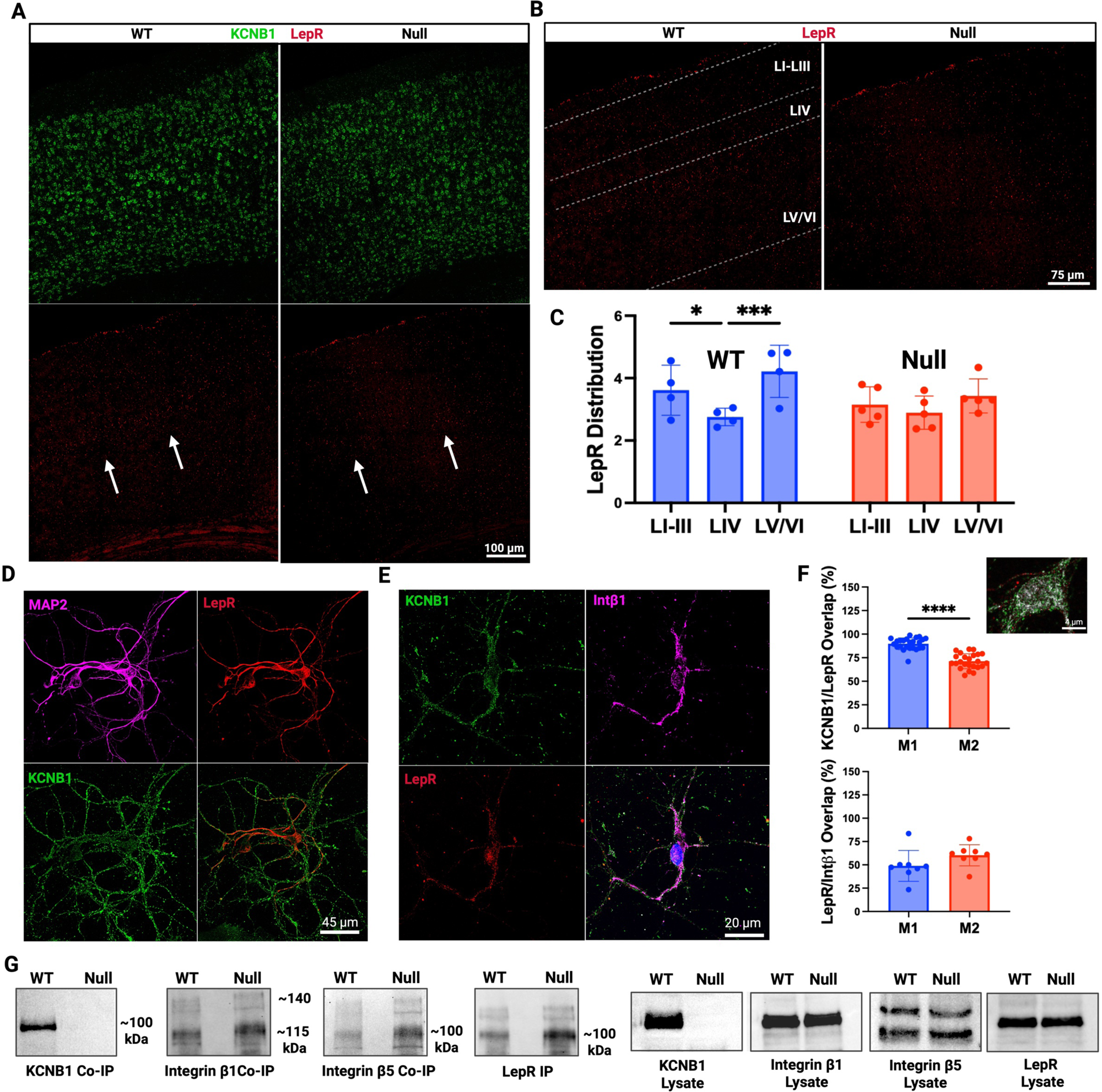
LepR forms cortical complexes with Kcnb1 and integrins. A) Representative confocal images of WT and Null cortical sections hybridized with fluorescent probes for Kcnb1 (blue) and LepR (green). B) Higher magnification of images in (A) highlighting the laminar distribution of LepR transcripts in WT cortex (arrows) which is disrupted in Null cortex. C) Quantifications of LepR transcripts segregated by cortical layers for WT (blue) and Null (red). N=2-3 brains/genotype with 4-5 technical replicates (points). D) Representative confocal images of primary cortical neurons co-immunostained for Kcnb1 (green), LepR (red), and Map2 (magenta), with merged overlays. E) Representative confocal images of primary cortical neurons co-immunostained for Kcnb1 (green), LepR (red), and integrin-β1 (magenta), with merged overlays. Note difference of scale with (D). F) M1 and M2 Pearson coefficients for Kcnb1/LepR overlap and LepR/integrin-β1 overlap. *Inset*: enlarged view of the images in (E) showing soma and basal dendrite co-localization of integrin-β1 and LepR immunofluorescence. G) Representative Western blots of LepR immunoprecipitates (IP) from cortical lysates, probed for Kcnb1 and integrin-β1 and integrin-β5 (co-IP). Input lysates showing total Kcnb1, integrin-β1, integrin-β5 and LepR are displayed on the right. All blots were obtained from the same antibody-stripped membrane.

### IKCs gate LepR signaling even without leptin

To probe IKC-mediated neurotrophic actions of leptin signaling, we blocked the leptin receptor with Allo-Aca (H-alloThr-Glu-Nva-Val-Ala-Leu-Ser-ArgAca-NH₂; Aca = 6-amino-caproic acid), a peptidomimetic competitive antagonist, and examined neuronal morphology and cytoskeletal dynamics (**Fig. 8**) (31). Remarkably, Allo-aca, alone or with leptin, reduced dendritic arborization in WT neurons, increased apical dendrite length, and did not alter the size of actin aggregates (**Fig. 8A**). In FRAP assays, Allo-aca (± leptin) raised the mobile fraction and accelerated recovery (k_off_), yielding WT responses that resembled those of Null neurons (**Fig. 8B**). In Null neurons, Allo-aca alone produced no detectable changes in morphology or actin dynamics. In contrast, combined with leptin it moderately improved over-arborization (SIP and apical dendrite length) while modestly enlarging actin aggregates (**Fig. 8C**). Moreover, the Allo-aca + leptin condition did not alter FRAP parameters in Null (**Fig. 8D**). Collectively, these data indicate that LepR can drive neurotrophic signaling in a ligand-independent mode gated by IKCs. Functionally, active LepR acts as a brake on the actin cytoskeleton, slowing actin polymerization. The residual normalizing effects of leptin on Null neuron morphology are consistent with this model. Since FRAP reflects acute applications, whereas the morphology assays involve chronic Allo-Aca exposure, incomplete LepR blockade can leave some residual leptin action (in fact, Allo-Aca targets site III but leaves the high-affinity site II available, limiting receptor activation rather than ligand binding). Accordingly, leptin’s effects under co-treatment are strongly attenuated relative to leptin alone (**Fig. 7**) but remain directionally consistent: dendritic complexity shifts toward normal and the apical dendrite shortens.

**Figure 8.**
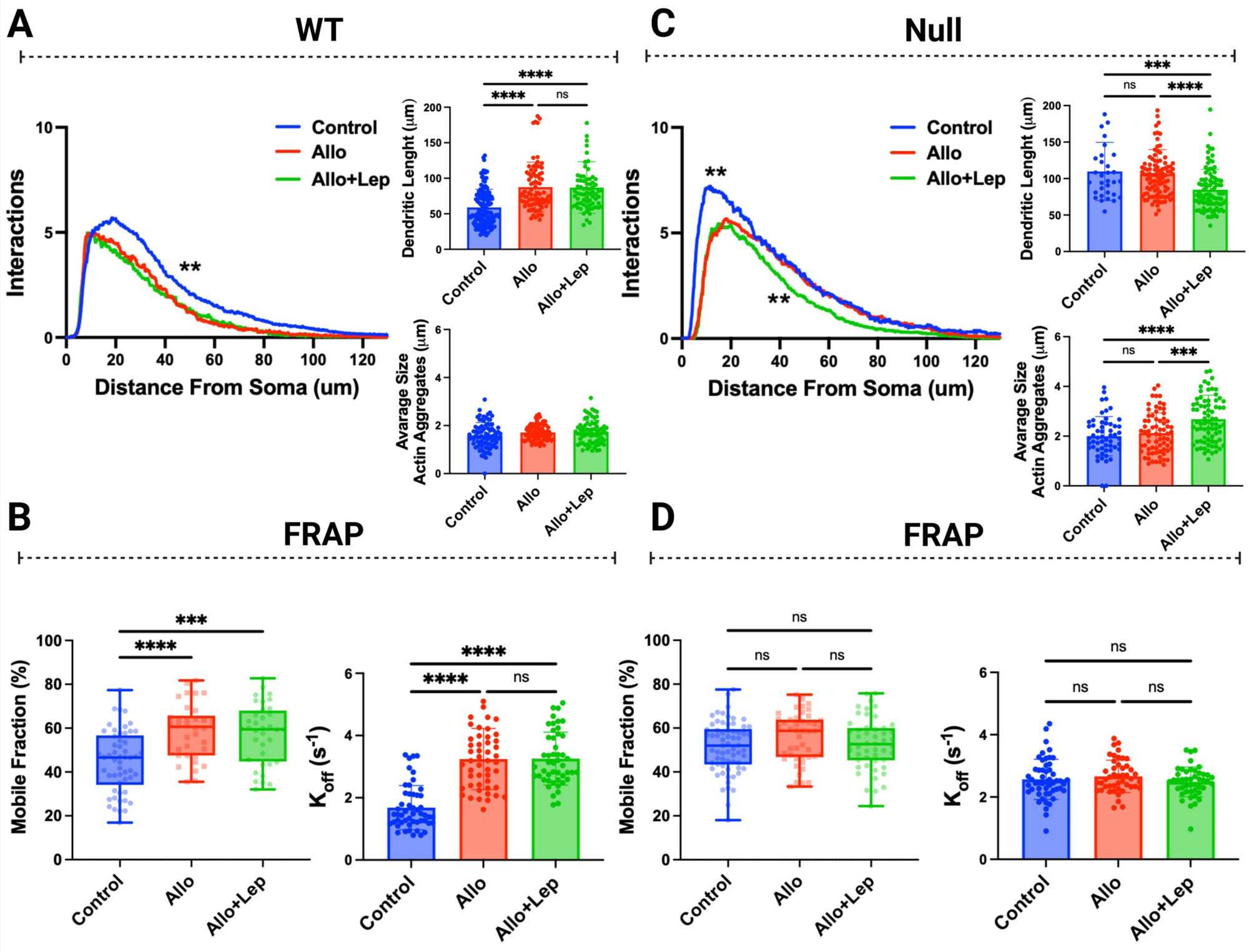
The IKC gates LepR in the absence of leptin. A) Mean SIP, apical dendrite length and average size of actin aggregates of WT neurons cultured under the indicated conditions. B) Mobile fraction (eqn. 4) and unbinding constant (k_off_, eqn. 3) of Lifeact-GFP–labeled actin in WT basal dendrites under the indicated conditions. C) Mean SIP, apical dendrite length and average size of actin aggregates of Null neurons cultured under the indicated conditions. D) Mobile fraction (eqn. 4) and unbinding constant (k_off_, eqn. 3) of Lifeact-GFP–labeled actin in Null basal dendrites under the indicated conditions. For morphological analyses, primary cortical neurons were cultured under control conditions or treated from DIV1 to DIV10 with leptin (0.3 μM, daily), Allo-aca (100 nM, every other day), or combinations of both. Neurons were fixed and immunostained for Map2 and actin at DIV10, imaged by confocal microscopy, and analyzed using Fiji. N=45-100 neurons genotype/condition from 3-7 biological replicates. For FRAP, neurons were transfected with Lifeact-GFP at DIV7, subjected to photobleaching at DIV8, and fluorescence recovery was quantified. Treatments were applied prior to imaging: Allo-aca (2 μM) or Allo-aca+leptin (1.3 μM) for 1 hour. N = 45-100 neurons per genotype/condition with 2 technical replicates from 3-5 biological replicates per genotype/condition. Statistical analyses included the Kolmogorov–Smirnov test for Sholl intersection profiles and one-way ANOVA with Tukey’s post hoc test for dendritic length and mobile fraction.

### Cortical neuron morphogenesis requires integrin signaling

Our co-immunoprecipitation and co-localization data raised a key mechanistic question: how does changing integrin activity influence leptin responsiveness? To test whether integrins can mediate leptin driven morphogenesis, we pharmacologically modulated RGD binding integrins using Cilengitide, an inhibitor of beta 1 and alpha 5 class RGD binding integrins, or fibronectin (FN), an extracellular matrix ligand that activates RGD binding integrins (32–34). The effects of these treatments, alone or in combination with leptin or Allo aca, on the Sholl intersection profile, apical dendrite length, actin aggregates, and the FRAP mobile fraction are summarized in **Fig. 9**. Fluorescence recovery curves and unbinding constants are shown in **Fig. S7**, and fluorescence intensity traces before, during, and after photobleaching are plotted in **Fig. S8**. Overall, these effects were internally consistent across assays, as already observed with leptin (**Fig. 6**) or Allo-aca (**Fig. 8**) alone. In agreement with prior work, in WT neurons fibronectin did not measurably alter the Sholl intersection profile, apical dendrite length, or actin polymerization, but it significantly reduced the mean size of actin aggregates (24). In contrast, Cilengitide impaired all readouts, supporting the idea that IKC associated integrins provide tonic signaling in WT cells. In Null neurons, both fibronectin and Cilengitide lowered the abnormally high rate of F-actin polymerization and, in doing so, ameliorated the associated morphological phenotypes. The effect of Cilengitide is readily explained if loss of Kcnb1 promotes integrin hyperactivation. Fibronectin could likewise act to correct aberrant integrin engagement in Null neurons, thereby restoring a more physiological activation state. In a context of integrin hyperactivation, this is compatible with a “biphasic clutch over engagement” mechanism, in which fibronectin can elicit a stop signal that suppresses actin dynamics (35–37). Notably, in both genotypes, LepR blockade or activation was dominant over the effects of integrin manipulation. This is evident when Allo-aca or leptin alone are compared with their respective combinations with fibronectin or Cilengitide across **Figs. 6**, **8** and **9** and **Figs S7B-F**. In summary, in WT neurons, integrin manipulation unmasked leptin susceptibility, and the minimal additivity between agents further supports the idea that LepR and integrins converge on a common pathway. Together, these results suggest a model in which integrins are the terminal effectors that couple actin polymerization to neuronal morphogenesis in WT neurons. Within this framework, both Kcnb1 and LepR regulate integrin signaling, as either LepR activation or inhibition or Kcnb1 ablation blunts integrin dependent control. The Null phenotypes are consistent with this scheme, since functionally disrupting the tripartite complex, through loss of Kcnb1 together with integrin perturbation and LepR blockade, abolishes the corrective effects of leptin, fibronectin, or Cilengitide. Thus, Kcnb1 and LepR modulate integrin activity to enable structural remodeling.

**Figure 9.**
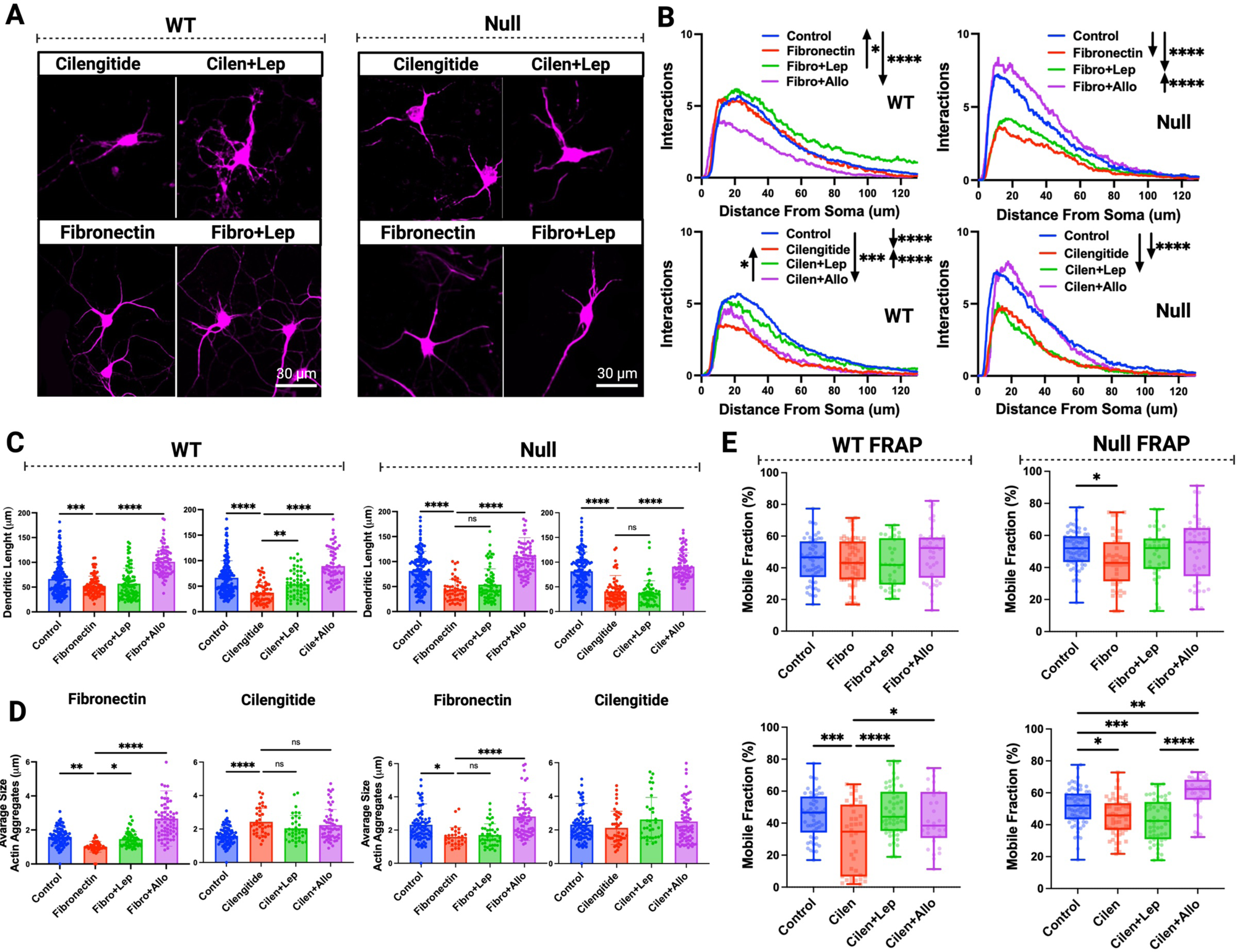
Leptin’s effects on cortical neurons require IKC activity. A) Representative confocal images of Map2-stained cortical neurons under the indicated conditions. B) Mean SIPs of WT and Null neurons cultured under the indicated conditions. C) Apical dendrite length of WT and Null neurons in the indicated conditions. D) Mean size of actin aggregates in WT and Null neurons in the indicated conditions. E) FRAP analysis of Lifeact-GFP–labeled actin in WT and Null basal dendrites under the indicated conditions. For morphological analyses primary cortical neurons were cultured under control conditions or treated from DIV1 to DIV10 with leptin (0.3 μM daily), fibronectin (50 μg/mL, coating), cilengitide (10 μM every other day), Allo-aca (100 nM, every other day) or combinations of leptin or Allo-aca with fibronectin or cilengitide. Neurons were fixed and immunostained for Map2 and actin at DIV10, imaged by confocal microscopy, and analyzed using Fiji. N = 45-100 neurons per genotype/condition from 3-7 biological replicates per genotype/condition. For FRAP, neurons were transfected with Lifeact-GFP at DIV7, subjected to photobleaching at DIV8, and fluorescence recovery was quantified. Treatments were applied prior to imaging: leptin (1.3 μM) and fibronectin (50 μg/mL) for 30 min; cilengitide (10 μM) or cilengitide + leptin for 2 h and Allo-aca (2 μM) alone or with fibronectin or cilengitide for 1 hour. N = 45-100 neurons per genotype/condition with 2 technical replicates from 3-5 biological replicates per genotype/condition. Statistical analyses included the Kolmogorov–Smirnov test for Sholl intersection profiles (SIPs) and one-way ANOVA with Tukey’s post hoc test for dendritic length and mobile fraction.

## DISCUSSION

To investigate how hormonal signaling regulates neurodevelopment, we focused on leptin in the Null mouse, which displays chronic hypoleptinemia due to hypothalamic dysfunction, and exhibits severe cortical malformations (17,18). We found that perinatal leptin supplementation during the first postnatal month markedly improved multiple aspects of the Null phenotype, including cognitive and sensorimotor behavior, neuronal migration, synaptic organization, brain anatomy, and neuronal morphology. These findings raise the possibility that disrupted leptin signaling contributes to neurodevelopmental deficits. However, in WT mice, altering leptin levels produced only marginal effects, despite robust rescue in Null animals. This juxtaposition poses a clear paradox: on the one hand, it points to a functional connection between a Kv channel and a hormone; on the other, it argues against simple hypoleptinemia as the principal cause of defective neurodevelopment, because WT should be highly leptin-responsive. Since Kcnb1 assembles with integrins into IKCs in cortex, we hypothesized that LepR resides in this complex and is able to modulate the neurodevelopmental activity of the IKC via its ligand (leptin). Consistent with this, co-immunoprecipitation and co-localization showed LepR associated with Kcnb1 and integrins in cortex, in line with hypothalamic observations (17). However, in primary cortical neurons, which are normally cultured without leptin, adding exogenous leptin produced only minimal adverse effects in WT cells, consistent with *in vivo* observations; by contrast, pharmacologic modulation of integrins sensitized WT neurons to leptin. In Null neurons, leptin normalized morphology and actin dynamics to WT levels, and critically, the effects of leptin and integrin modulation were non-additive, further indicating that they operate along the same IKC–LepR axis rather than in parallel pathways. Together, these findings indicate that leptin’s modulation of LepR is transduced through the IKC to regulate actin-cytoskeletal remodeling (38–43). This raises a second paradox: if leptin acts through the IKC, why are its effects small in WT? We propose that IKCs impose a tonic constraint on LepR, limiting additional ligand drive. Indeed, blocking LepR in WT neurons, whether or not exogenous leptin was present, accelerated filament growth and induced morphological defects that mirror the Null phenotype, whereas the same manipulations in Null neurons had minor impact. This pattern suggests LepR carries tonic activity even without leptin, leading us to propose that LepR is subject to dual control by leptin and the IKC (**Fig. S9**). In this context, leptin’s normalization of Null neurons most likely reflects restoration of IKC-dependent pathways that are normally masked in WT. Consistent with this model, the exaggerated dendritic arborization and elevated actin polymerization in Null versus WT indicate that, under physiological conditions, the IKC–LepR complex functions as a brake on both processes.

Despite these advances, several important questions remain. Rodents experience a postnatal leptin surge peaking between P9 and P16, with both timing and magnitude dependent on nutrition (44). Accordingly, WT pups nursed by Null dams failed to undergo a surge comparable to that of normally weaned WT animals. However, since WT mice are normoleptinemic, it is difficult to disentangle the consequences of missing the surge from overall leptin sufficiency. A definitive test will require a hypothalamus-specific conditional Kcnb1 knockout or a constitutively leptin-deficient model such as the ob/ob mouse (26). Although actin remodeling emerges as a major downstream target of IKC–LepR signaling, the precise molecular mechanism is still unclear. How does LepR physically engage IKCs to influence cytoskeletal dynamics? Do integrins act in a correlative manner or are they essential mediators of this process? Our findings point to a complex interplay among Kcnb1, integrins, and LepR, with Kcnb1 serving as a central hub for leptin–integrin interactions. Further work will be needed to delineate the structural and signaling logic of this tri-partite complex. Additional players may also contribute, for example, specific capping proteins could facilitate IKC–LepR interactions while simultaneously shielding LepR from direct leptin binding. Importantly, our analysis did not address the role of glia, cells fundamental to neuronal development and very likely involved in regulating these processes (45,46). Alternative models are conceivable as well. One possibility is that leptin and the IKC act independently on shared neurodevelopmental pathways, with the IKC normally exerting dominance over leptin.

However, this scenario seems unlikely, since it would have to apply across the diverse processes involved, including migration, dendritogenesis, synaptogenesis, and cognitive function. These mechanistic uncertainties acquire particular importance in the clinical context. Given the role of KCNB1 in DEE and the occurrence of not only missense but also nonsense mutations in this gene (typically heterozygous), in affected children, it is intriguing to consider that leptin supplementation might offer therapeutic benefit (47).

Beyond the neurodevelopmental abnormalities examined here, this might extend to seizure susceptibility, a hallmark feature of both DEE patients and our DEE mouse models, including the Null, but not directly assessed in the present study (18,20).

In summary, our results uncover a previously unrecognized pathophysiological framework linking ion channel signaling, cytoskeletal remodeling, and metabolic hormones. Defining this interplay will not only advance our understanding of Kv channel biology in brain development, but may also open therapeutic opportunities for conditions such as DEE and maternal obesity, all of which are known to compromise neurocognitive outcomes in early life (6–9,11).

## METHODS

### Table of reagents

| REAGENT or RESOURCE | SOURCE | IDENTIFIER |
| --- | --- | --- |
| **Antibodies** |  |  |
| MAP2 Polyclonal Antibody | Thermo Fisher Scientific | Cat# PA1-10005<br>RRID:AB_1076848 |
| Synapsin 1 Monoclonal Antibody (7H10G6) | Thermo Fisher Scientific | Cat# MA5-31919<br>RRID:AB_2787542 |
| BrdU Monoclonal Antibody (BU-1) | Thermo Fisher Scientific | Cat# MA3-071<br>RRID:AB_10986341 |
| PSD-95 Polyclonal Antibody | Thermo Fisher Scientific | Cat# 51-6900<br>RRID:AB_2533914 |
| Anti-Neural cell adhesion molecule L1 Antibody, clone 324 | Millipore Sigma | Cat# MAB5272<br>RRID:AB_2133200 |
| Anti-Actin Antibody, clone C4 | Millipore Sigma | Cat# MAB1501<br>RRID:AB_2223041 |
| Mouse Anti-Neurofilament, Medium, 160 kD Monoclonal Antibody, Unconjugated, Clone NF-09 | Abcam | Cat# AB7794<br>RRID:AB_306083 |
| Anti-BrdU antibody [BU1/75 (ICR1)] | Abcam | Cat# AB6326<br>RRID:AB_305426 |
| Anti-Kv2.1/KCNB1 antibody [K89/34] | Abcam | Cat# AB192761<br>RRID:AB_3677394 |
| Mouse Leptin R Antibody | R&D Systems | Cat# AF497<br>RRID:AB_2281270 |
| Mouse Leptin R Biotinylated Antibody | R&D Systems | Cat# BAF497<br>RRID:AB_2296953 |
| Integrin β-5 (D24A5) Rabbit mAb | Cell Signaling Technology | Cat# 3629<br>RRID:AB_2249358 |
| Integrin β-1 Antibody | Cell Signaling Technology | Cat #4706<br>RRID:AB_823544 |
| Goat anti-Chicken IgY (H+L) Cross-Adsorbed Secondary Antibody, Alexa Fluor™ Plus 647 | Thermo Fisher Scientific | Cat# A32933<br>RRID:AB_2762845 |
| Donkey anti-Mouse IgG (H+L) Highly Cross-Adsorbed Secondary Antibody, Alexa Fluor™ Plus 488 | ThermoFisher Scientific | Cat# A32766<br>RRID:AB_2762823 |
| Goat anti-Mouse IgG (H+L) Cross-Adsorbed Secondary Antibody, Alexa Fluor™ Plus 594 | ThermoFisher Scientific | Cat# A11005<br>RRID:AB_2534073 |
| Donkey anti-Rabbit IgG (H+L) Highly Cross-Adsorbed Secondary Antibody, Alexa Fluor™ Plus 488 | ThermoFisher Scientific | Cat# A32790<br>RRID:AB_2762833 |
| Goat anti-Rabbit IgG (H+L) Highly Cross-Adsorbed Secondary Antibody, Alexa Fluor™ 594 | ThermoFisher Scientific | Cat# A11037<br>RRID:AB_2534095 |
| Donkey anti-Rabbit IgG (H+L) Highly Cross-Adsorbed Secondary Antibody, Alexa Fluor™ 647 | ThermoFisher Scientific | Cat#A-31573 |
| Goat anti-Rat IgG (H+L) Cross-Adsorbed Secondary Antibody, Alexa Fluor™ 594 | ThermoFisher Scientific | Cat# A11007<br>RRID:AB_10561522 |
| Donkey anti-Goat IgG (H+L) Highly Cross-Adsorbed Secondary Antibody, Alexa Fluor™ Plus 594 | ThermoFisher Scientific | Cat# A32758 |
| Rabbit anti-Mouse IgG (H+L) Secondary Antibody, HRP | ThermoFisher Scientific | Cat# 61-6520 |
| Goat anti-Rabbit IgG (H+L) Secondary Antibody, HRP | ThermoFisher Scientific | Cat# 31460 |
| Rabbit anti-Goat IgG (H+L) Secondary Antibody, HRP | ThermoFisher Scientific | Cat# 31402<br>RRID:AB_228395 |
| **Chemicals** |  |  |
| Cilengitide | MedChemExpress | Cat# HY-16141 |
| Allo-aca | MedChemExpress | Cat# HY-P3212 |
| Fibronectin Bovine Plasma | Millipore Sigma | Cat# F1141 |
| Sodium Tetraborate Decahydrate | Millipore Sigma | Cat# B9876 |
| Sucrose | Millipore Sigma | Cat# S0389 |
| Normal Donkey Serum, Sterile | Millipore Sigma | Cat# 566460 |
| Donkey Serum | Millipore Sigma | Cat# D9663 |
| Bovine Serum Albumin | Millipore Sigma | Cat# A9418 |
| Protein G-Agarose | Millipore Sigma | Cat#<br>11719416001 |
| Dulbecco's Phosphate Buffered Saline | Millipore Sigma | Cat# D1408 |
| Triton™ X-100 | Millipore Sigma | Cat# T8787 |
| Hydrochloric acid 37%, ACS reagent | Millipore Sigma | Cat# 258148-2.5L |
| Mouse Leptin Recombinant Protein, PeproTech® | ThermoFisher Scientific - Gibco™ | Cat# 450-31-1MG |
| BrdU (5-Bromo-2'-Deoxyuridine) | ThermoFisher Scientific | Cat# B23151 |
| Streptavidin Agarose Resin | ThermoFisher Scientific | Cat# 20349 |
| UltraPure™ Agarose | ThermoFisher Scientific | Cat# 16500500 |
| Hydrochloric acid, 36.5-38.0%, NF, Multi-Compendial, GMP, J.T.Baker™ | ThermoFisher Scientific - Avantor™ | Cat# 9544-33 |
| RNAscope® Probe- Mus musculus leptin receptor (Ob-r) mRNA complete cds | ACD, (Biotechne) | Cat# 402731-C2 |
| RNAscope® Probe - Mm-Kcnb1-C3 | ACD, (Biotechne) | Cat# 316941-C3 |
| Tissue-Tek, O.C.T Compound | Sakura | Cat# 4583 |
| Protease inhibitor Cocktail Set I | Calbiochem | Cat# 539131 |
| 30% Acrylamide/Bis Solution | Bio-rad | Cat# 1610158 |
| Trans-Blot Turbo 5x Transfer Buffer, 1L | Bio-rad | Cat# 10026938 |
| TEMED | Bio-rad | Cat# 1610800 |
| **Media and Reagents** |  |  |
| Precoated coverslips GG-15-1.5-PDL | Neuvitro | Cat# GG-15-PLL |
| Precoated coverslips GG-12-1.5-PDL | Neuvitro | Cat# GG-12-1.5-PDL |
| Fisherbrand™ Superfrost™ Plus Microscope Slides Precleaned Clipped Corners | Fischer Scientific | Cat# 22-037-246 |
| Non-Fat dry milk bioKeMIX 500GM | LabScientific | Cat# M0841 |
| Opti-MEM™ I Reduced Serum Medium | ThermoFisher<br>Scientific - Gibco™ | Cat# 31985062 |
| Lipofectamine™ 2000 Transfection Reagent | ThermoFisher<br>Scientific | Cat# 11668019 |
| B-27™ Supplement (50X), serum free | ThermoFisher<br>Scientific - Gibco™ | Cat# 17504044 |
| L-Glutamine (200 mM) | ThermoFisher<br>Scientific - Gibco™ | Cat# 25030081 |
| Penicillin-Streptomycin (10,000 U/mL) | ThermoFisher<br>Scientific - Gibco™ | Cat# 15140122 |
| Fetal Bovine Serum | Millipore Sigma | Cat# F4135 |
| Sodium Pyruvate (100 mM) | ThermoFisher<br>Scientific - Gibco™ | Cat# 11360070 |
| Neuronal Culture Medium for Pierce™ Primary Cell<br>Isolation Kits | ThermoFisher<br>Scientific | Cat# 88283 |
| Basal Medium Eagle (BME) | ThermoFisher<br>Scientific - Gibco™ | Cat# 21010046 |
| Fluoromount-G™ Mounting Medium, with DAPI | ThermoFisher<br>Scientific -<br>Invitrogen™ | Cat# 00-4959-52 |
| HBSS 1X Hanks' Balanced Salt Solution | ThermoFisher<br>Scientific - Gibco™ | Cat# 14025-092 |

| **Commercial assays** |  |  |
| --- | --- | --- |
| Leptin Mouse ELISA Kit | ThermoFisher Scientific | Cat# KMC2281 |
| Neural Tissue Dissociation Kit (P) | ThermoFisher Scientific | Cat# 130-092-628 |
| Pierce™ BCA Protein Assay Kits | ThermoFisher Scientific | Cat# 23225 |
| SuperSignal™ West Pico PLUS Chemiluminescent Substrate | ThermoFisher Scientific | Cat# 34580 |
| Rapid GolgiStain™ Kit | FD NEUROTECHNOLOGIES, INC. | Cat# PK401A |
| RNAscope® Intro Pack for Multiplex Fluorescent Reagent Kit | ACD, (Biotechne) | Cat# 323280 |
| **Organisms/strains** |  |  |
| Mouse: C57BL6/J background | The Jackson Laboratory | Strain #:000664<br>RRID:IMSR_JAX:000664 |
| Mouse: NULL mouse | Genome Editing Share Resource Core Facility at Rutgers | C57BL/6J-Kcnb1em2Sesf/J |
| **Microscopes** |  |  |
| Zeiss Axiophot | Zeiss |  |
| Olympus FV1000MPE | Olympus |  |
| Nikon Eclipse Ti2 serie | Nikon |  |
| Leica TCS SP8 tauSTED 3X (SP8tauSTED3X) | Leica |  |
| Wide Field Leica DMI8 Thunder System | Leica |  |
| **Software and algorithms** |  |  |
| Neuroanatomy Fiji ImageJ | Tiago Ferreira | https://imagej.net/update-sites/neuroanatomy/ |
| EthoVision XT | N/A | https://www.noldus.com/ethovision-xt |
| Colocalization Threshold Fiji ImageJ (Coloc 2) | Tony Collins | https://imagej.net/plugins/colocalization-threshold |
| Prism | GraphPad by Dotmatics | https://www.graphpad.com/scientific-software/prism/ |
| ANY-maze- Behavior and Video Tracking Software |  | https://www.any-maze.com |
| Image J | NIH | https://imagej.net/software/fiji/ |

### Animals

All procedures involving animals were approved by the Rutgers University Institutional Animal Care and Use Committee (IACUC; protocol #PROTO999900293), and all experiments were conducted in accordance with relevant ethical guidelines for animal research. Wild-type (C57BL/6J) and homozygous knock-in (KI) mice of either sex were used at two developmental stages: embryonic day 13 (E13), postnatal 7 (P7) and 4 and 8 weeks of age. The KI line used is C57BL/6J-Kcnb1^em2Sesf/J^ (referred to as Null), which harbors a premature stop codon after serine 336, producing a truncated Kcnb1 protein that is rapidly degraded and fails to reach the plasma membrane (18). Littermates of both sexes were randomly assigned to experimental groups.

### Leptin Treatments

Lyophilized recombinant leptin was reconstituted in sterile 20 mM Tris-HCl (pH 8.0) to a stock concentration of 1.0 mg/mL, following the manufacturer’s instructions.

Intraperitoneal (i.p.) administration. The stock solution was freshly diluted in 0.9% saline to a final volume of 0.5 µg/g in a final volume of 100 µL for adults. For postnatal pups from P0-P14 the injections were performed subcutaneously in the back and from P15-P28 intraperitoneal with a final volume of 50 µL.

*In vitro* neuronal treatments. Leptin was added every morning to the neuronal media reaching a concentration of 0.3 µM from DIV (Day In Vitro) 1 to DIV10.

FRAP (Fluorescence Recovery After Photobleaching). Leptin was added 30 minutes prior the live imaging at a final concentration of 1.3 µM.

### Hormone Measurements

Submandibular blood samples were collected 90 minutes after intraperitoneal injection or 24-hour post-injection from 4-week-old mice and after behavioral experiments in 8-week-old mice. After collection, samples were left at room temperature for 20 minutes and centrifuged at 4 °C at 14,000 RPM for 20 minutes. The serum was collected and analyzed via ELISA according to the manufacturer’s instructions and quantified using an Infinite M Nano+ plate reader (Tecan, Männedorf, Switzerland).

### Behavioral protocols

#### Open Field Test

The open field test was conducted following the standard protocol described by Seibenhener et al. (48). Briefly, testing took place under dim lighting in a dedicated behavior room containing a plexiglass arena (50×50×40 cm) with walls lined with tissue paper to block external visual stimuli. A video tracking system (EthoVision XT; Noldus Information Technology, Leesburg, VA) was used to record mouse movement. Each mouse was habituated to the arena 24 hour prior the experiment for 5 minutes. The day of the experiment the animals acclimated to the testing room for 30 minutes. The animal was then placed in the center of the arena, and the experimenter exited the room. Locomotor activity was recorded for a single 10-minute session.

#### Nest Building Test

Mice were individually housed in clean, enrichment-free cages and provided with a pre-weighed nestlet pad as nesting material. After 16 hours, the nest was photographed. Nests were scored on a scale from 1 to 5 according to established nest-building criteria (49).

#### Marble Burying Test

Each mouse was placed in a clean standard cage filled with 10 cm of fresh bedding. Twenty glass marbles (14 mm diameter) were arranged evenly across the surface. After 30 minutes, the mouse was removed, and the cage was photographed. Three blinded evaluators counted the number of marbles buried at least two-thirds deep in bedding. Following each trial, the marbles and cage were cleaned with 70% ethanol, and the bedding was discarded.

#### Y-maze

The Y-maze consisted of three identical arms (35 cm long × 10 cm wide × 15 cm high) arranged at 120° angles. Mice were individually placed at the center of the arms and allowed to freely explore the maze for 5 min in a quiet behavioral room under uniform lighting (∼100 lux). Arm entries were recorded by a digital overhead camera. An entry was defined as the placement of all four paws into an arm. Between animals, the maze was thoroughly cleaned with 70% ethanol to eliminate olfactory cues.

Spontaneous alternations were scored manually from video recordings by an observer blinded to treatment. A correct alternation was defined as consecutive entries into all three arms without repetition (e.g., ABC, BCA), an incorrect alternation was defined when the animal repeated arms visits.

#### Barnes maze

The Barnes maze consisted of a circular platform 100 cm in diameter, elevated 75 cm above the floor, and made of opaque material. Twenty evenly spaced holes (5 cm diameter) were positioned around the perimeter, but only one led to an escape box (target). The escape box contained fresh bedding, which was replaced between animals. The platform was divided into four equal zones of five holes each: the “Target zone” (containing the escape hole), the “Positive zone” (right of Target zone), the ”Negative zone” (left of Target zone), and the “Opposite zone” (opposite to Target zone). The maze surface and escape box were cleaned with 70% ethanol between animals.

##### Habituation (Day 0)

Mice were placed in the center of the maze and allowed to explore for 2 min. During habituation, the escape box was located in the Negative zone. If a mouse did not locate the escape box within 2 min, it was gently guided to the box and allowed to remain there for 1 min.

##### Acquisition (Training)

Twenty-four hours after habituation, mice underwent two training trials (Trials 1–2) with the escape box fixed in the Target zone. Each trial began with the mouse placed at the center of the maze and allowed up to 3 min to locate the escape box. If the mouse did not enter the box within the allotted time, it was guided to it and allowed to remain inside for at least 30 s. Forty-eight hours later, mice underwent three additional training trials (Trials 3–5) under the same conditions. Two time points were recorded, the time to locate and enter the escape box. These five trials were used to assess learning.

##### Probe Test

Seventy-two hours after Trial 5, a probe trial was performed to assess spatial memory. The escape box was removed, leaving all holes visually identical. The mouse was placed in the maze center and allowed to explore freely for 90 s. The location of the (now absent) escape hole was unchanged. Behavior was recorded with a digital overhead camera and analyzed using ANY-maze software. Outcome measures included time spent in each zone, time to first approach of the target hole, and number of visits/pokes to the target versus other holes.

#### Novel object recognition

##### Training (familiarization)

Two identical golf balls were placed in opposite corners of the arena, ∼6 cm away from the walls. Objects were heavy enough to prevent displacement by the animals. Each mouse was placed in the center of the arena and allowed to explore freely for 5 min. Following the training session, animals were returned to their home cages for a 24 hours retention interval.

##### Test

Twenty-four hours after training, one of the two familiar objects was replaced with a novel object (a Lego cube) of comparable size but different shape and texture. Mice were again placed in the center of the arena and allowed to explore freely for 5 min. Behavior was recorded with an overhead video tracking system (EthoVision XT, Noldus Information Technology, Leesburg, VA) and analyzed using ANY-maze software. Object exploration was defined as directing the snout toward the object within 2 cm and/or touching/sniffing the object.

##### Data analysis

The discrimination index (DI) was calculated as:

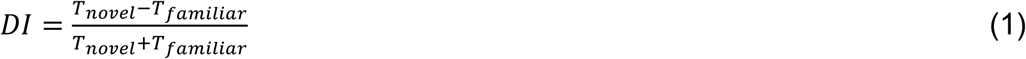

where *T_novel_* is the time spent exploring the novel object and *T_familiar_* is the time spent exploring the familiar object. A positive DI indicates preference for the novel object.

Total exploration time was also analyzed to ensure no group differences in exploratory activity. Animals that explored both objects for <5 s in total during the test were excluded from analysis. The test was performed in the same arena of the Open Field.

### Cell Cultures

#### Primary Cortical Neurons

Primary neuronal cultures were prepared from embryonic day 13 (E13) mouse cortices. Pregnant dams were euthanized by CO₂ inhalation followed by cervical dislocation in accordance with institutional animal care guidelines. Under a dissection microscope, cortical tissue was isolated in ice-cold sterile Hank’s Balanced Salt Solution (HBSS). Tissue was mechanically dissociated using fine tweezers and pooled into a 15 mL conical tube. Excess HBSS was removed by aspiration. Enzyme Mix 1 (Enzyme P + Buffer X; Neural Tissue Dissociation Kit (P), Miltenyi Biotec, Cat. No. 130-092-628) was added, and the sample was incubated for 15 min at 37 °C in a water bath. After incubation, Enzyme Mix 2 (Enzyme A + Buffer Y, same kit) was added. The tissue was gently triturated 10–15 times with a P1000 pipette and incubated for an additional 10 min at 37 °C. Following incubation, tissue was further dissociated by gentle pipetting (10–15 times). Ten mL of plating medium was added [composition per 20 mL: 18.31 mL Basal Medium Eagle (with Earle’s salts), 1 mL heat-inactivated fetal bovine serum (FBS), 200 μL sodium pyruvate (100× stock), 200 μL L-glutamine (200 nM final), and 100 μL penicillin/streptomycin (200×)]. The suspension was centrifuged at 300g for 10 min at room temperature. The supernatant was aspirated and cells were resuspended in 1 mL plating medium. After counting with a Neubauer chamber, neurons were plated onto 15 mm poly-D-lysine–coated coverslips (Neuvitro, GG-12-1.5-PDL) at a density of ∼150,000 cells per dish in 1.5 mL plating medium.

Cultures were maintained at 37 °C in a humidified incubator with 5% CO₂. After 24 h, plating medium was replaced with Neurobasal medium supplemented with B27 (Thermo Fisher Scientific).

##### In Vitro Assays

All *in vitro* experiments were conducted blind to genotype.

Pharmacological agents were freshly diluted from stock solutions to the indicated final concentrations and added directly to the culture medium. Primary cortical neurons were treated with leptin (0.3 μM), fibronectin (50 μg/mL, coating), cilengitide (10 μM), Allo-Aca (100 nM), or combinations thereof. Treatments were initiated at DIV1 and continued until DIV10, when neurons were fixed for immunocytochemistry. Leptin was freshly supplemented daily whereas Allo-Aca and cilengitide were added every other day.

Fibronectin was applied as a coating substrate at a final concentration of 5 μg/cm². Briefly, wells containing 15 mm poly-D-lysine–coated coverslips were covered with 1 mL of fibronectin solution (diluted in sterile water) for 30 s then the cover slips were let air-dry at room temperature for 45 min under sterile conditions before plating neurons.

##### Neuronal Staining

At DIV10, neurons were washed once with 1X phosphate-buffered saline (PBS) for 5 min, then fixed with a 50:50 acetone:methanol solution on ice for 20 min. Cells were washed three times for 5 min each with PBS. Neurons were blocked with 3% bovine serum albumin (BSA) in PBS for 1 hour at room temperature. Primary antibodies (diluted 1:500–1:800 in blocking solution) were applied overnight at 4°C. After three PBS washes (5 min each), appropriate fluorophore-conjugated secondary antibodies (diluted 1:500–1:800) were applied for 1 hour at room temperature, followed by three additional PBS washes. Coverslips were mounted with Fluoromount-G™ containing DAPI (Thermo Fisher) and stored at 4°C. Staining was visualized and images were taken with a Nikon CREST X-Light LFOV confocal microscope.

#### Fluorescence Recovery After Photobleaching (FRAP)

Primary neurons derived from E13 embryos were cultured to DIV7. Neurons were transfected using Lipofectamine with 1.5 μg of LifeAct-GFP plasmid and 5 μL of transfection reagent per well of a 12-well plate and incubated for 16 h. Neurons were either treated or immediately analyzed after the incubation time. Treatments were applied prior to imaging: leptin (1.3 μM) and fibronectin (50 μg/mL) for 30 min, Allo-aca (2 μM) or Allo-aca+leptin for 1 h; cilengitide (10 μM) or cilengitide + leptin for 2 h. FRAP experiments were performed on an Olympus FV1000MPE confocal microscope. Imaging was carried out at 2 Hz (1 frame every 0.5 s) for a total of 24 frames. Three pre-bleach frames were acquired as baseline, followed by photobleaching for 1.5 s. The first post-bleach frame was recorded at 2 s. Fluorescence recovery was measured in basal dendrites ∼5–15 μm from the soma, where Kcnb1 expression is enriched. Photobleaching parameters were kept constant across all experiments, and control conditions were included in each session. FRAP analysis was done with ImageJ. The fluorescence in the area of bleach was measured from frame 1 to frame 24 and the measurement was normalized over the background fluorescence. Fluorescence recovery, R(t), was expressed as a percentage:

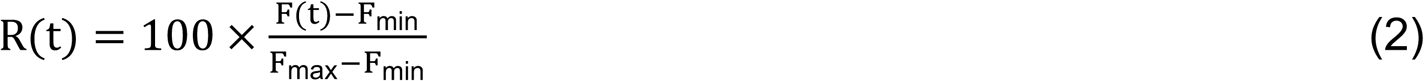

and fitted to a single exponential function:

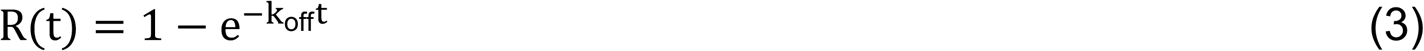

where k_off_ is the rate of unbinding of LifeAct-GFP from F-actin. The mobile fraction was calculated as:

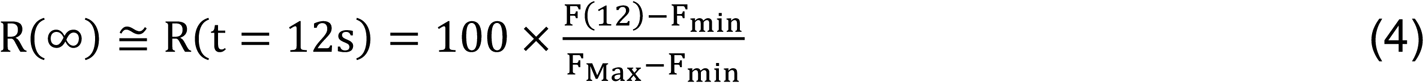

#### Neurons analysis

Single-neuron morphology was quantified using the ImageJ *Sholl Analysis* plugin. Concentric circles were drawn from the soma center with a starting radius of 0.275 µm, continuing in 0.275 µm increments up to a final radius of 130 µm. The Sholl decay was calculated using a log–log transformation, and both linear and normalized interaction plots were generated. For dendritic length measurements, the longest visible dendrite for each neuron was recorded. Spine density and morphology were calculated using the SpineJ plugin in ImageJ. Actin-rich puncta were quantified with the ImageJ *Analyze Particles* function. Images were first thresholded, followed by background separation using the *Process → Binary → Watershed* command. Regions of interest (ROI) were then selected, and particles were analyzed within a size range of 0.5–100 µm² and a circularity range of 0–1.

### Biochemistry

All biochemical experiments, including Western blotting and immunofluorescence, were conducted blind to genotype.

#### Crude Brain Lysates

Animals were euthanized via CO₂ asphyxiation followed by cervical dislocation. Brains were rapidly extracted, and cortices were dissected on ice. Tissue was transferred to 1.7 mL conical tubes and homogenized using a plastic pestle in lysis buffer containing 0.32 M sucrose, 5 mM Tris-HCl (pH 6.8), 0.5 mM EDTA, supplemented with protease inhibitor cocktail and 0.1% Triton X-100 (Sigma-Aldrich).

Homogenates were further processed by sonication in an ice-cooled water bath for 1–3 min (e.g., 10-s pulses with 10-s rests). Samples were kept on ice throughout the procedure. Homogenates were centrifuged at 14,000 × g for 10 min at 4°C, and the supernatants were collected and stored at −80°C for subsequent analysis.

#### Immunoprecipitation

To reduce nonspecific binding, lysates were pre-cleared with pre-washed streptavidin beads for 30 min at room temperature. Following centrifugation at 5,000 × g for 1 min, the supernatants were incubated overnight at 4°C with the primary antibody. Streptavidin beads were blocked with 3% bovine serum albumin (BSA) in PBS, added to the antibody–lysate mixtures, and incubated for 8 hours at 4°C.

Beads were then washed four times with PBS, resuspended in 4× SDS sample buffer (10% SDS, 30% glycerol, 0.5 M Tris-HCl, 0.02% bromophenol blue) containing 2% β-mercaptoethanol, and heated at 95–100°C for 5 min. After centrifugation at 10,000 × g for 1 min, the supernatants were collected and analyzed by SDS-PAGE followed by Western blotting.

#### Western Blotting

Protein concentration (∼60 µg per sample) was determined using the Bradford assay. Samples were prepared in 5X SDS sample buffer containing 5% β-mercaptoethanol, heated at 95–100 °C for 5 min, and centrifuged at 10,000g for 1 min. Proteins were separated on 10–15% SDS-PAGE gels, transferred onto nitrocellulose membranes, and blocked with 5% nonfat dry milk in TBST (TBS + 0.1% Tween-20) for 1 h at room temperature. Membranes were incubated overnight at 4 °C with primary antibodies (1:1000), followed by horseradish peroxidase (HRP)-conjugated secondary antibodies (1:2000) for 1 h at room temperature. Protein bands were detected using SuperSignal™ West Pico PLUS chemiluminescent substrate and visualized on a Bio-Rad ChemiDoc imaging system. Actin was used as a loading control.

#### Membrane Stripping

Nitrocellulose membranes were stripped for reuse by incubating twice for 10 min in Mild Stripping Buffer (1.5% glycine, 0.1% SDS, 1% Tween 20, pH 2.2), followed by PBS washes (2×10 min) and TBST washes (2×5 min). Membranes were re-blocked and re-probed as described above.

### Tissue Processing and Imaging

#### Neonatal Brains and BrdU

Pregnant dams at embryonic day 16.5 (E16.5) received intraperitoneal injections of 5-bromodeoxyuridine (BrdU) at 100 mg/kg body weight from a freshly prepared 10 mg/mL stock in sterile PBS. Separately, dams were injected with either saline (control) or 0.5 mg/kg body weight leptin until birth of the litter. Then offsprings were subcutaneously injected from postnatal day 0 (P0) to P7 with 0.5 mg/kg body weight leptin every other day. At P7, pups were anesthetized with ketamine/xylazine, perfused with saline, followed by 4% paraformaldehyde (PFA).

Brains were post-fixed in 4% PFA overnight at 4°C and cryoprotected in 30% sucrose. Brains were embedded in agarose and sectioned at 70 µm throughout the cortex using a vibratome (Leica VT1000S). For antigen retrieval, sections were incubated in 2.0 N HCl for 30 min in a water bath at 37°C. Brain slices were then washed with 1X PBS, incubated with 0.1M Borax Solution 8.0 pH (sodium tetraborate decahydrate) for 10 minutes twice and finally washed again with 1X PBS. Sections were incubated in blocking/permeabilization buffer (donkey serum in PBS with 0.4% Triton X-100, pH 7.43), followed by overnight incubation at 4°C with BrdU primary antibody. After washing, appropriate secondary antibodies were applied for 2 hours at room temperature, followed by additional PBS washes. sections were mounted with Fluoromount-G™ containing DAPI (Thermo Fisher), coverslipped and sealed.

#### Adult Brains

Mice were anesthetized with ketamine/xylazine (100 mg/kg and 10 mg/kg, i.p.) and perfused with saline followed by 4% PFA at room temperature. Brains were post-fixed in 4% PFA overnight at 4 °C, rinsed in PBS, and cryoprotected in a sucrose gradient (10%, 20%, 30%) at 4 °C until tissues sank. Brains were frozen in Optimal Cutting Temperature (OCT) compound with isopentane pre-cooled to –20 °C and stored on dry ice. Coronal sections (40 μm) were cut. Slices were then blocked with 5% Donkey Serum in PBS with 0.3% Triton X-100. Then, they were incubated with primary antibodies in 3% Donkey Serum over night at 4 °C, washed in PBS, and incubated with fluorophore-conjugated secondary antibodies for 1 hour at room temperature. After additional PBS washes, sections were mounted with Fluoromount-G™ containing DAPI (Thermo Fisher), coverslipped, and sealed. Slides were imaged and analyzed with Leica TCS SP8 tauSTED 3X super-resolution microscope.

#### Golgi Staining

Golgi staining was performed using the FD Rapid GolgiStain™ Kit (FD NeuroTechnologies, Columbia, MD), according to the manufacturer’s protocol. Brains were sectioned at 100 µm thickness, mounted on gelatin-coated slides and stained with Golgi-staining solution. Images were acquired using a Widefield Leica DMI8 Thunder System Microscope and a Zeiss Axio Imager M1 in 5x or 63x. Sholl analysis and Spine analysis was performed using ImageJ/Fiji software.

#### RNA scope

After rinsing with saline, brains were perfused with 4% PFA. They were then removed and fixed overnight in 4% PFA. Tissue was cryoprotected by immersion in increasing concentrations of sucrose (10%, 20%, and 30%) before embedding in OCT compound and freezing at −80 °C. Coronal sections (10 μm, one or two per slide) were prepared on a cryostat and processed the next day for *in situ* hybridization following the protocol provided by the manufacturer (ACD, Bio-Techne, Minneapolis, MN). The *Kcnb1* probes were designed to detect a 20-base pair sequence in the channel’s C-terminal region; in the null allele, the stop codon occurs in the exon encoding the S4 segment. Imaging was carried out on a Leica TCS SP8 tauSTED 3X super-resolution system, and images were analyzed using FIJI software.

#### Statistics and Reproducibility

Sample size (N) was estimated via power analysis using α = 0.05 and power = 0.8 (50). Data were assessed for normality and log-normality using the D’Agostino and Pearson omnibus test. Statistical significance was defined as P < 0.05. Statistical comparisons were performed using one-way ANOVA followed by Tukey’s post hoc test, Brown-Forsythe and Welch ANOVA with Dunnett’s T3 multiple comparisons, two-tailed paired and unpaired Student’s *t*-test, or the two-sample Kolmogorov-Smirnov test, as appropriate. Analyses were conducted using GraphPad Prism software. Data are presented as mean ± standard error of the mean (SEM).

## Data availability

All data generated or analyzed in this study, including the source data underlying the figures, have been deposited in Dryad and will be made publicly available upon publication of this manuscript. For inquiries prior to that time, please contact the corresponding author.

## Acknowledgements

This study was supported by an NSF grant (#2030348) to FS.

## Authors contribution

EF designed experiments, performed experiments analyzed data; Anika G performed experiments and analyzed data; Aniket G performed Golgi and behavioral analysis and analyzed data, MdCM performed behavioral analysis, primary neurons and analyzed data; RP performed biochemistry and analyzed data; SA maintained mice colonies and analyzed data, CI performed immunofluorescence and analyzed data; KL analyzed data; SV analyzed data and FS directed the study and wrote the manuscript.

