## supplemental figures S1-S9 for "The IKC-leptin axis functions as a molecular bridge between neurodevelopment and endocrine signaling"

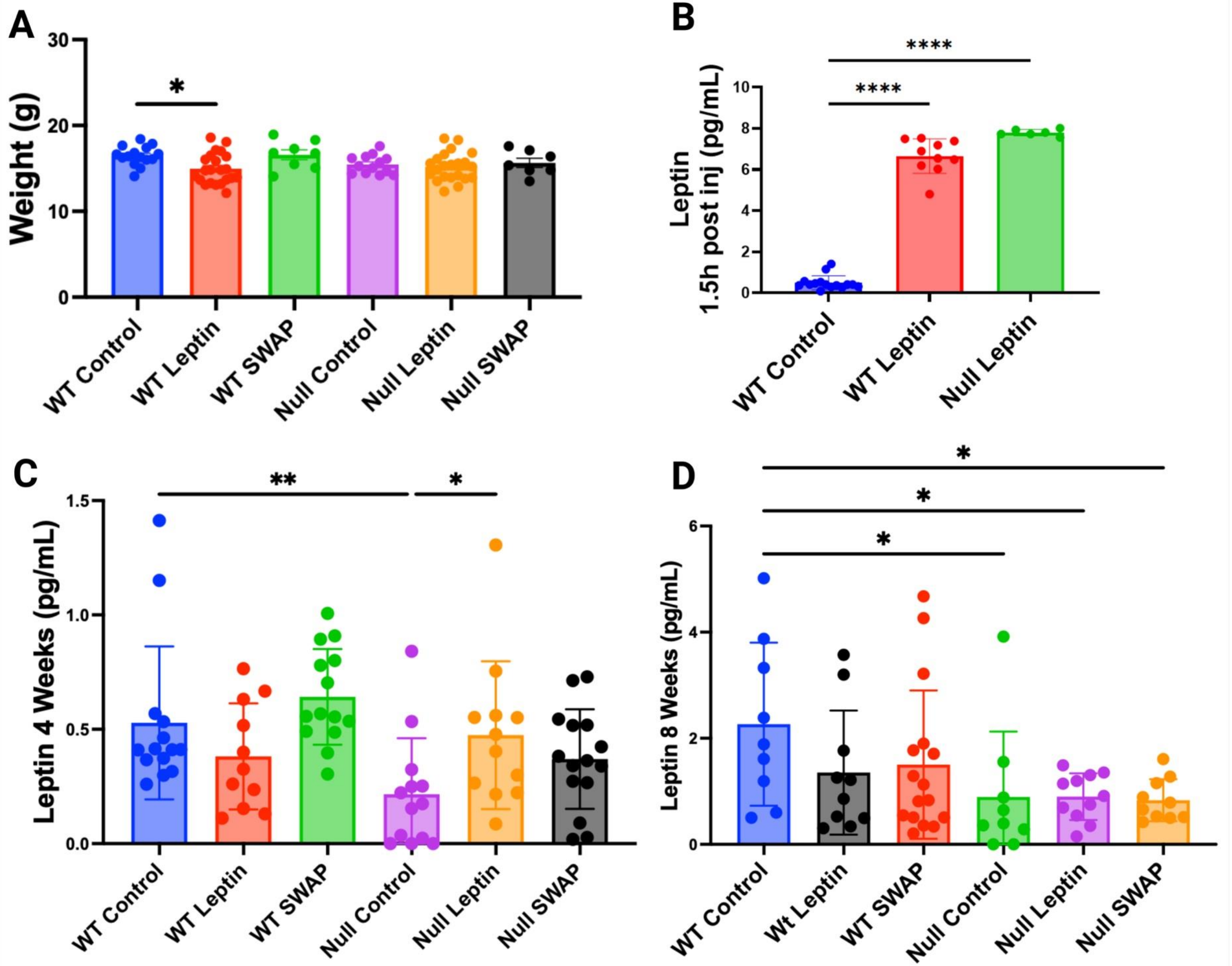


**Fig. S1 Effects of leptin treatment on body weight and circulating leptin levels**

(A) Mean body weight of 4-week-old WT and Null mice under the indicated conditions.
(B) Serum leptin 90 min after an injection of recombinant leptin in 4-week-old WT and Null mice; saline-injected WT control shown for comparison.
(C) Basal circulating leptin in 4-week-old mice by genotype/condition.
(D) Basal circulating leptin in 8-week-old mice by genotype/condition.
Four-week-old mice received daily saline (control), recombinant leptin (0.5 mg/kg, P0–P28), or were cross-fostered (SWAP). Body weight was measured at 4 weeks; circulating leptin was measured at 4 and 8 weeks. N=8-15 mice per genotype/condition. Statistical comparisons used one-way ANOVA with Tukey's post hoc test.

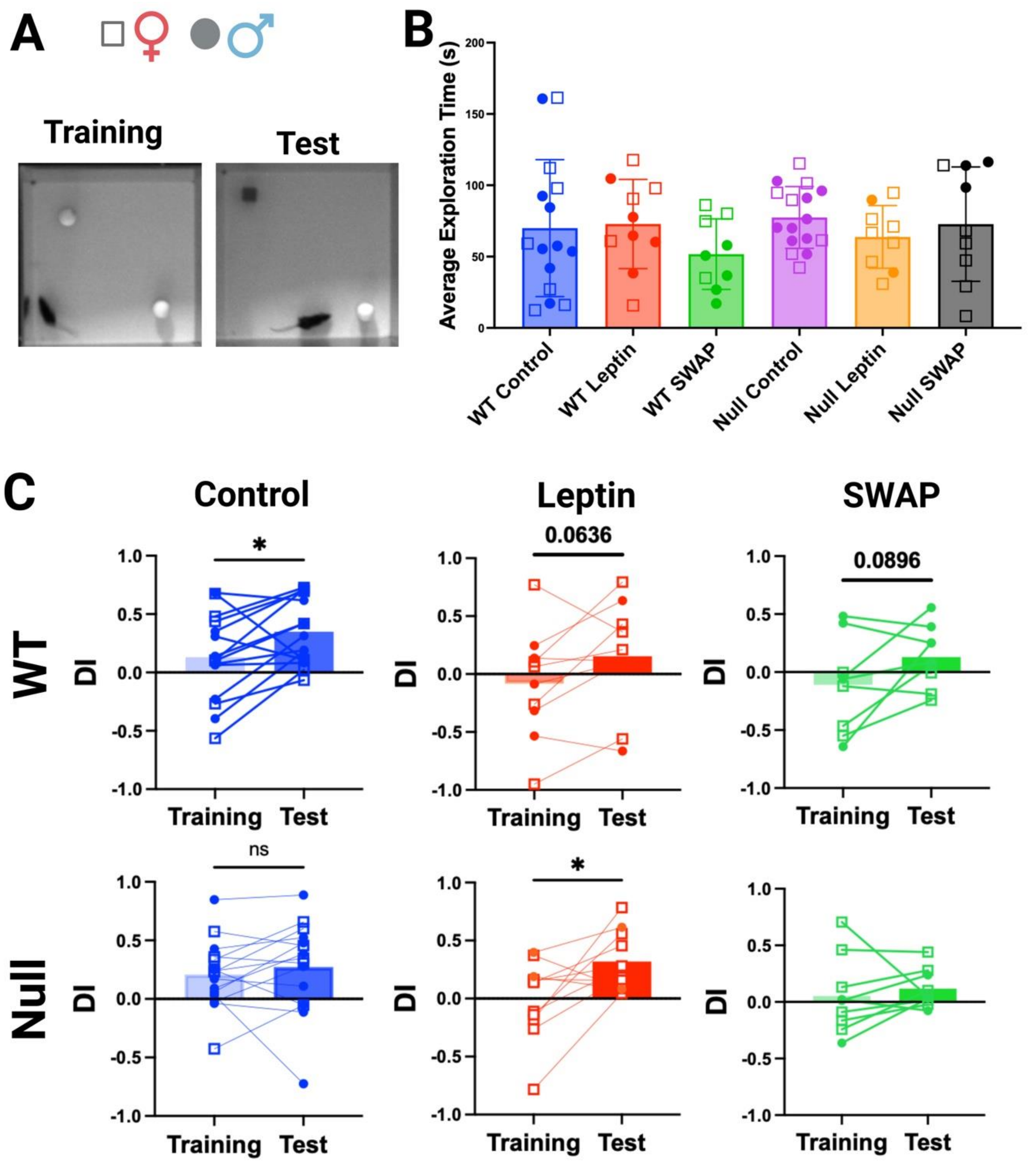


**Fig. S2 Leptin and cross-fostering rescue Novel Object Recognition in Null mice**

A) Representative images of a WT mouse during the sample (training) phase with two identical familiar objects (golf balls) and during the test phase with one novel object (Lego cube).

B) Mean exploration time by genotype/condition across phases (average of total time exploring the objects during training and test).

C) Discrimination Index (DI; eqn. 1) trajectories over time for each genotype/condition.

“Leptin”: daily recombinant leptin (0.5 mg/kg, P0–P28). “SWAP”: cross-fostered pups. Statistics: one-way ANOVA with Tukey’s post hoc test. “.

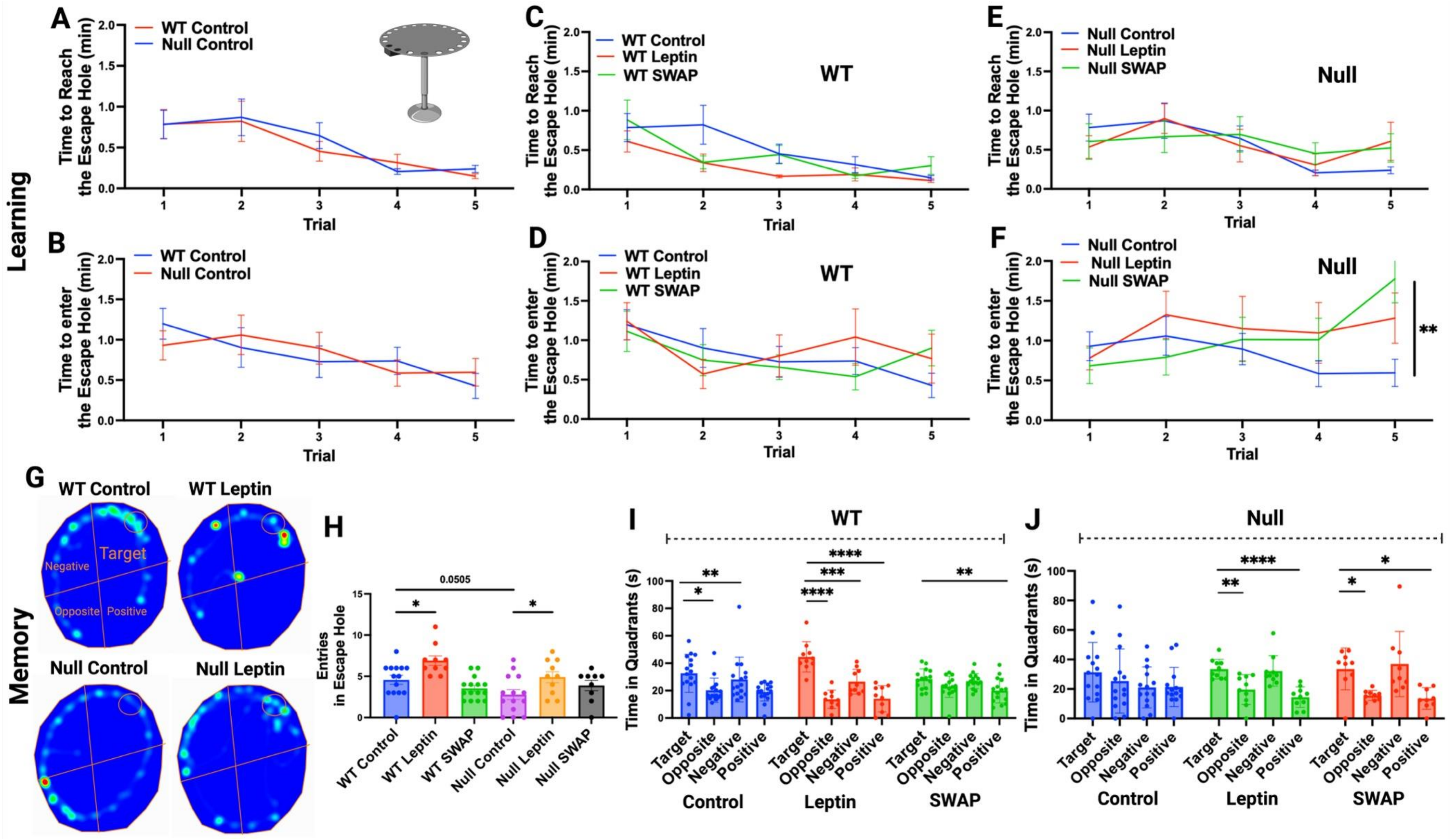


**Fig. S3 Leptin supplementation and cross-fostering improve Barnes maze recall in WT and Null mice**

A–B) Time to locate (A) and to enter (B) the escape hole during acquisition (training days 1–5) in WT and Null mice.

C–D) Time to locate (C) and to enter (D) the escape hole during acquisition for WT mice reared under the indicated conditions.

E–F) Same measures as in (C–D) for Null mice.

G) Representative heat maps from the recall (probe) phase showing search behavior of WT and Null mice under the indicated rearing conditions. The four quadrants and the escape hole location are marked.

H) Mean number of entries into the escape hole during the recall phase for the indicated genotypes/conditions.

I–J) Time spent in each quadrant during the recall phase for WT (I) and Null (J) mice under the indicated rearing conditions.

“Leptin”: daily recombinant leptin (0.5 mg/kg, P0–P28). “SWAP”: cross-fostered pups. Statistics: one-way ANOVA with Tukey’s post hoc test for multiple comparisons.

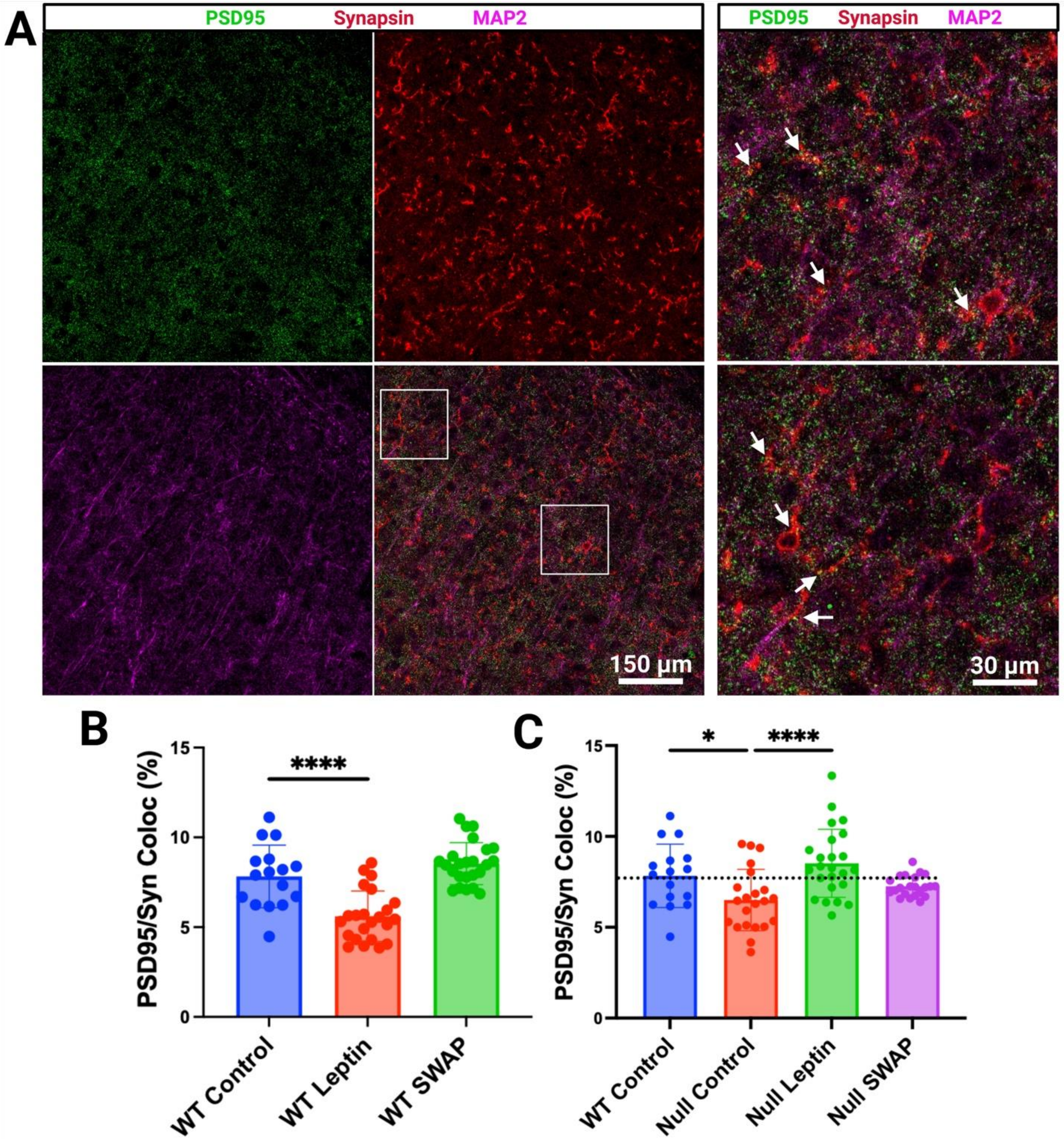


**Fig. S4 Leptin and cross-fostering enhance PSD95–Synapsin-1 co-localization in Null cortex**

A) Representative confocal images of PSD95 (postsynaptic) and Synapsin-1 (presynaptic) and Map2 immunofluorescence in cortical sections from WT mice. Right panels show magnified views of boxed regions on the left; arrows indicate PSD95–Synapsin-1 co-localized puncta (putative synapses).
B) Quantification of PSD95–Synapsin-1 co-localization in WT under the indicated rearing/treatment conditions.
C) Quantification of PSD95–Synapsin-1 co-localization in Null under the indicated conditions, with WT control shown for comparison.
"Leptin": daily recombinant leptin (0.5 mg/kg, P0–P28). "SWAP": cross-fostered pups. N=3 brains per genotype/condition with N=8 technical replicates. Statistics: one-way ANOVA with Tukey's post hoc test. Co-localization was computed in Fiji using the image calculator and analyze particles tools.

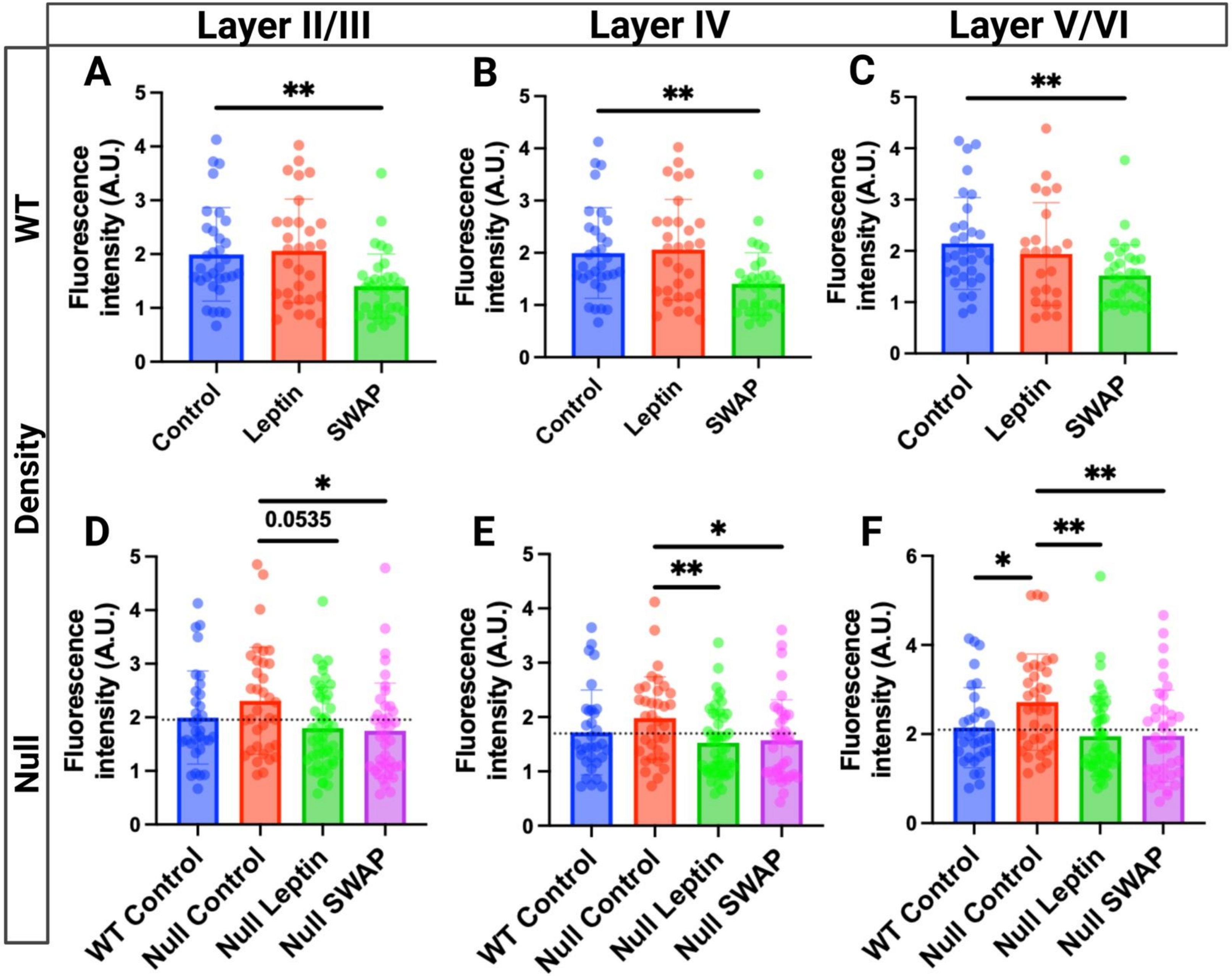


**Fig. S5 Leptin normalized NFM fluorescence in the Null**

A-C) NFM fluorescence in the indicated cortical layers (A: LII/III), (B: LIV), (C: LV/VI) in brain slices of WT reared under the indicated conditions.
D-F) As in (A-C) for Null slices.
Eight-week-old male and female mice received daily injections of saline (control), or 0.5 mg/body weight recombinant leptin (leptin) from P0–P28, or were cross-fostered (SWAP), as described elsewhere. N=3-5 brains/condition with 5-7 technical replicates/brain (points in graphs). Statistical comparisons were performed using one-way ANOVA with Tukey’s post hoc test.

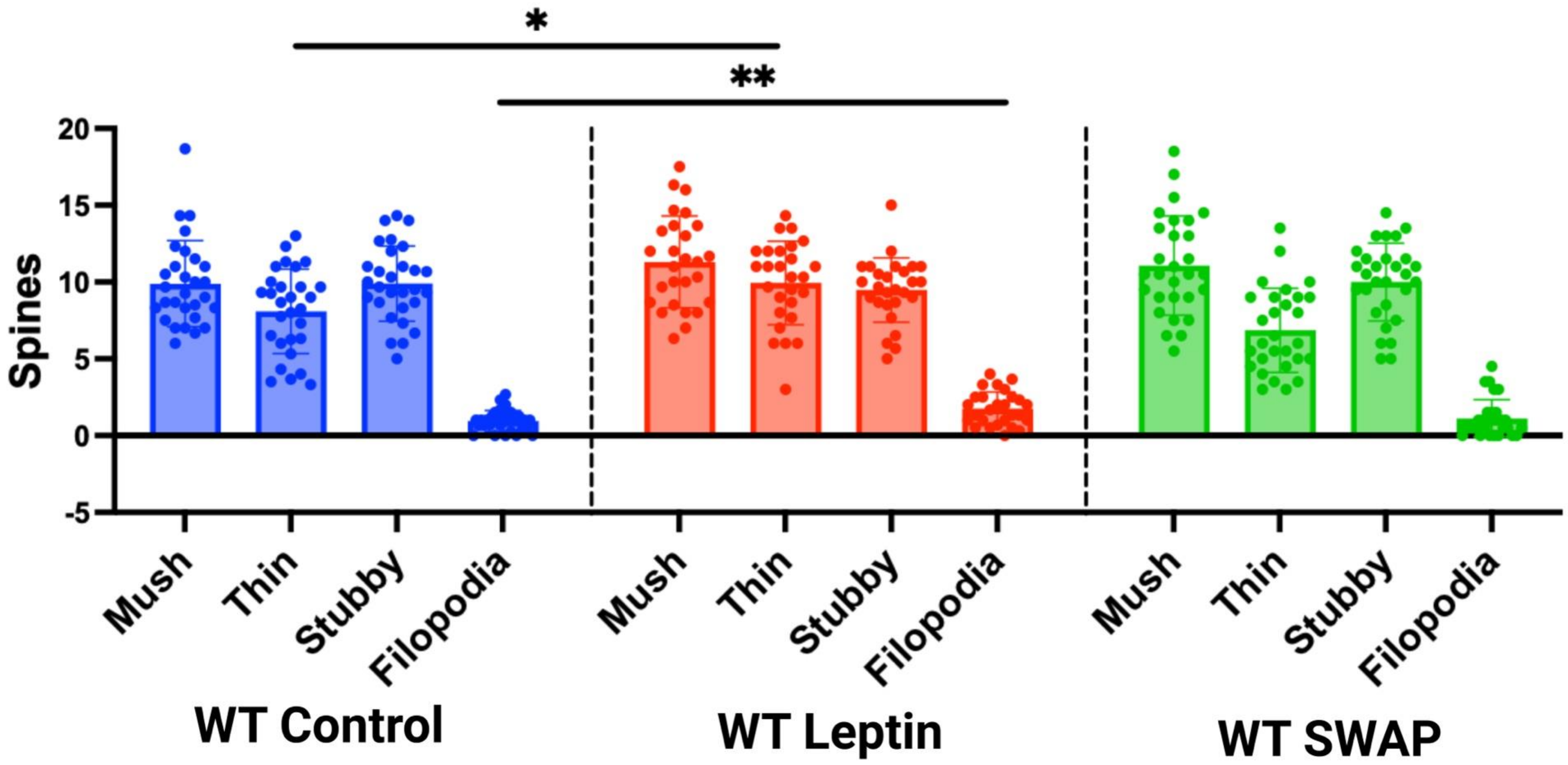


**Fig. S6 Leptin delays spine maturation in WT**

Mean counts of WT dendritic spines by morphological class across the indicated conditions. Four-week-old mice received daily saline (control), recombinant leptin (0.5 mg/kg, P0–P28), or were cross-fostered (SWAP). Brains were processed with Golgi staining and imaged by confocal microscopy (N = 3 brains/condition; 8–10 images/brain). Spines were counted along fixed dendritic segments (45 µm; 3 dendrites/image) and averaged across individual data points. Statistical comparisons used one-way ANOVA with Tukey's post hoc test. Spines were counted and analyzed using SpineJ plugin in ImageJ.

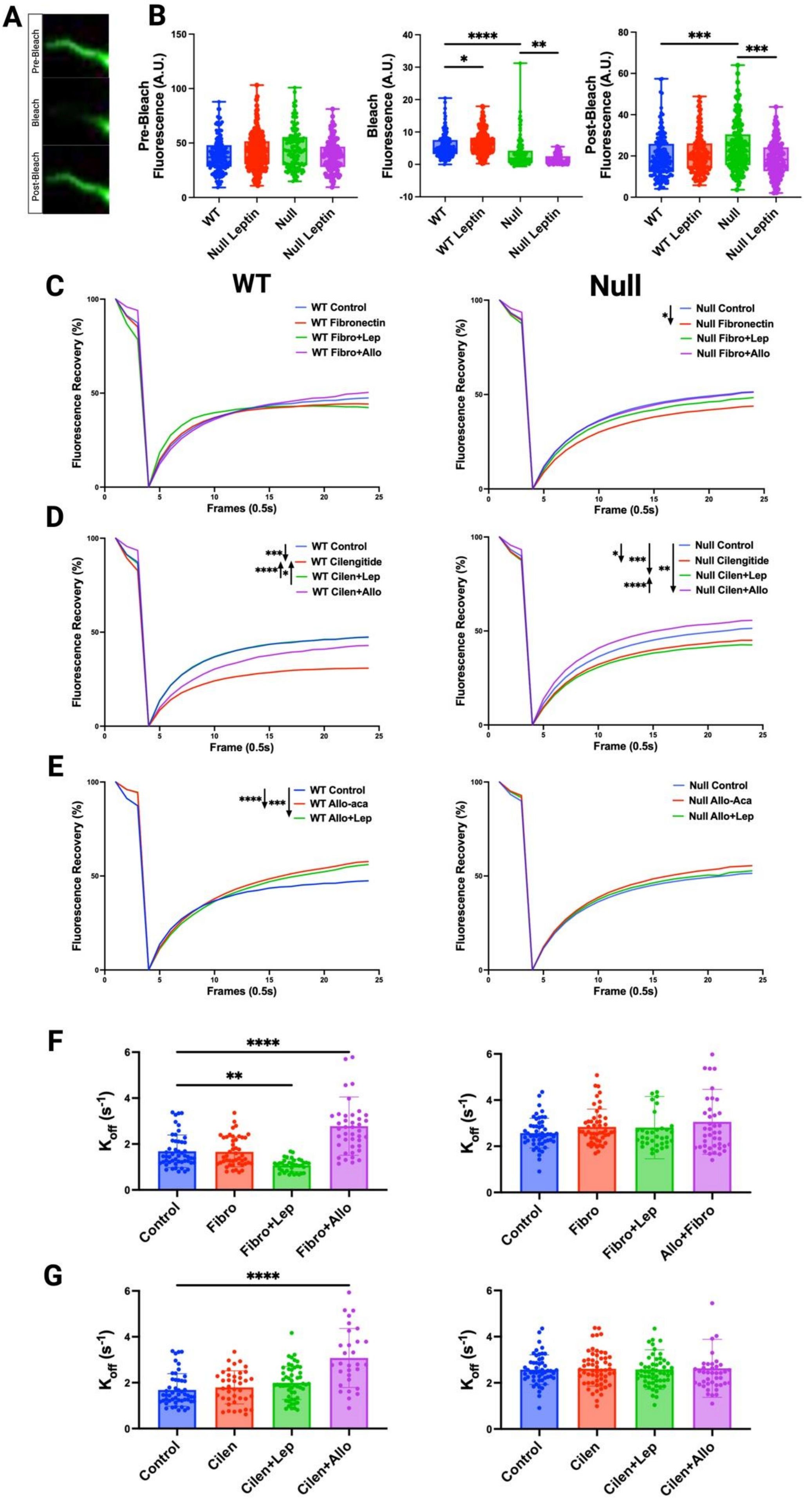


**Fig. S7 FRAP analysis of dendritic actin with LepR and integrin ligands/antagonists**

A) Representative Lifeact-GFP fluorescence in a WT basal dendrite before (pre-bleach, Frame 1) during (bleach, Frame 4) and 12 s after (post-bleach, Frame 24).

B) Fluorescence intensity quantification before, during, and after photobleaching for WT and Null under the indicated conditions.

C-E) Fluorescence recovery curves for WT and Null under the indicated conditions. SEMs are not shown for clarity.

F-G) $K_{off}$ constants (eqn. 3) for WT and Null under the indicated conditions.

Primary cortical neurons were transfected with Lifeact-GFP at DIV7, subjected to photobleaching at DIV8 and fluorescence recovery was quantified. 30 minutes prior the experiment neurons were incubated with leptin (1.3 μM), fibronectin (50 μg/mL), and 2 hours with cilengitide (10 μM) and Allo-aca (2 μM), or in combinations. N = 45-100 neurons per genotype/condition with 2 technical replicates from 3-5 biological replicates per genotype/condition . Statistical analyses were calculated using one-way ANOVA with Tukey's post hoc test.

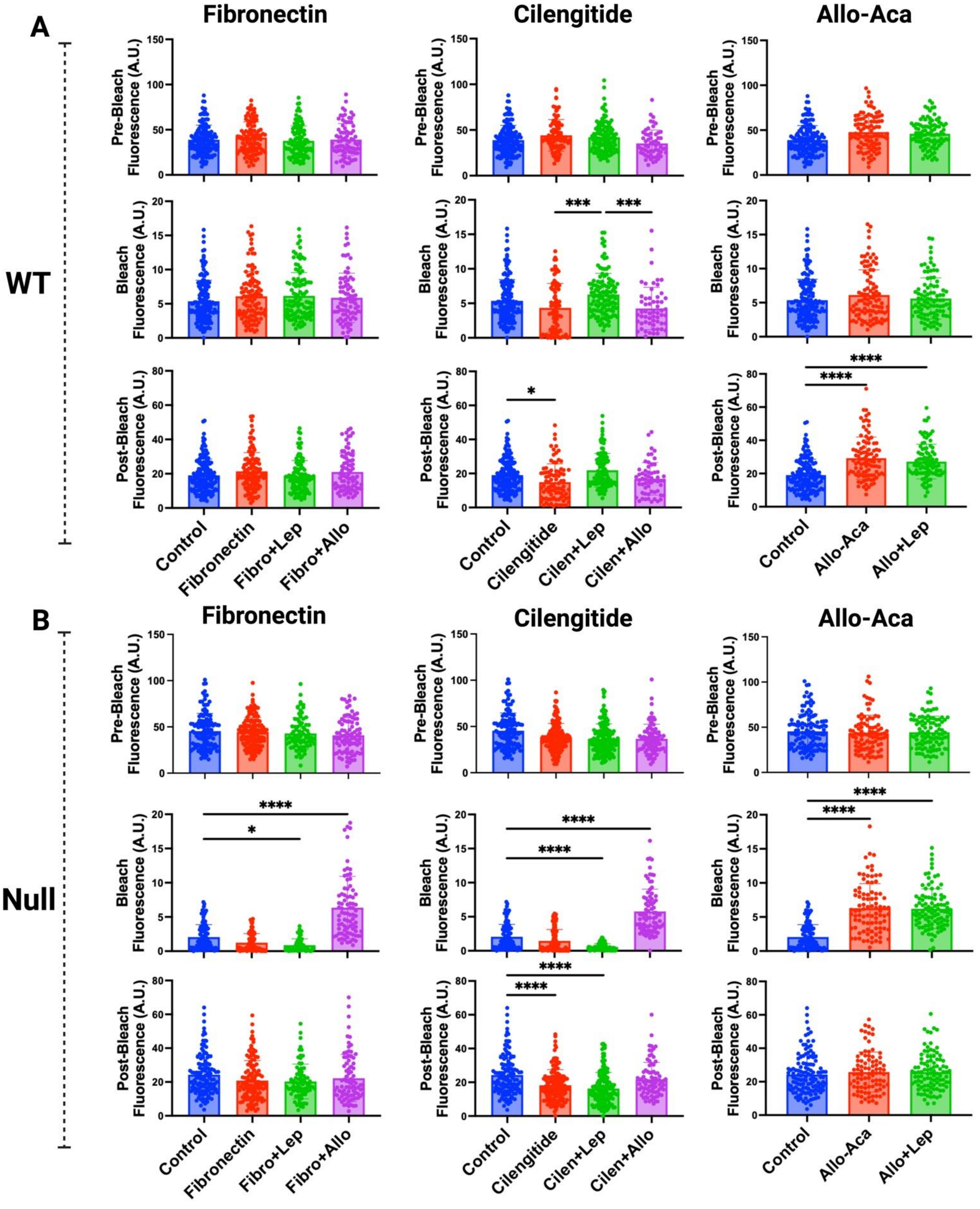


**Fig. S8 | Fluorescence intensity traces in WT and Null neurons before, during, and after photobleaching with LepR and integrin ligands/antagonists.**

A–B) Quantification of fluorescence intensity before, during, and after photobleaching in WT (A) and Null (B) neurons under the indicated conditions.

Primary cortical neurons were transfected with Lifeact-GFP at DIV7, photobleached at DIV8, and fluorescence recovery was quantified. Neurons were treated 30 min before imaging with leptin (1.3 μM) and/or fibronectin (50 μg/mL), and 2 h before imaging with cilengitide (10 μM) and/or Allo-aca (2 μM), alone or in combination. N = 45–100 neurons per genotype/condition, with 2 technical replicates from 3–5 biological replicates per genotype/condition. Statistical analysis was performed using one-way ANOVA with Tukey's post hoc test.

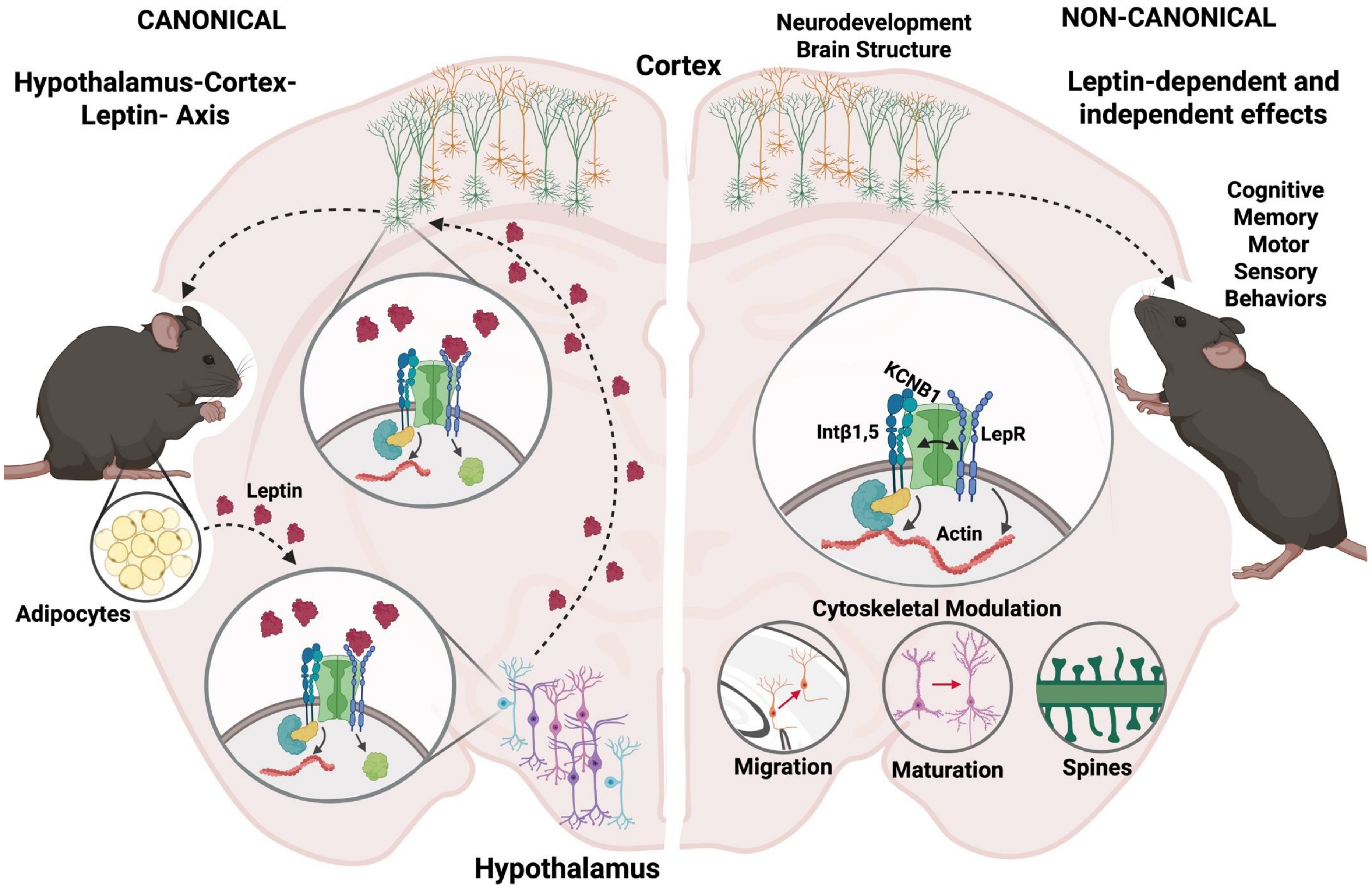


**Fig. S8. Dual-mode model of LepR–IKC neurotrophic signaling.** Tripartite complexes comprising the leptin receptor (LepR) and integrin–Kv channel complexes (IKCs) form a feedback loop with circulating leptin: IKC activity in hypothalamic $ARH^{POMC}$ neurons helps set leptin levels, while leptin shapes brain-wide neurodevelopment. In the canonical mode, leptin binds to LepR to engage IKC-dependent developmental pathways. In the non-canonical mode, IKCs gate LepR directly. In both modes, LepR activity, transduced via integrins, acts as a brake on actin polymerization and thereby restrains dendritogenesis and spinogenesis. In IKC-deficient states (e.g., Null), chronic hypoleptinemia and impaired IKC function converge to drive neurodevelopmental abnormalities, consistent with both arms of the model.
